# Mesenchymal Stromal Cells regulate human Hematopoietic Stem Cell survival, engraftment and regeneration via PGE2/cAMP signaling pathway

**DOI:** 10.1101/2023.11.09.566361

**Authors:** Siva Sai Naga Anurag Muddineni, Chen Katz Even, Adi Zipin-Roitman, Diana Rasoulouniriana, Debanjan Singha Roy, Rabeaa Aborgies, Katia Beider, Yael Raz, Neta Solomon, Gal Hershkovitz, Yael Shulman, Rony Chen, Eviatar Weizman, Claudia Waskow, Arnon Nagler, Michael Milyavsky

## Abstract

Ionizing radiation (IR) and chemotherapies severely impair hematopoietic stem and progenitor cell (HSPC) function, causing bone marrow failure and secondary malignancies. Mesenchymal stromal cells (MSCs) within the hematopoietic niche support HSPC survival and regeneration, but the underlying pro-survival mechanisms remain incompletely understood. Here, we show that MSCs suppress IR-induced apoptosis in human HSPCs and preserve their regenerative capacity. Transcriptomic analyses identified a robust induction of CREB target genes in HSPCs upon MSC contact, driven by MSC-secreted prostaglandin E2 (PGE2) via cAMP signaling. While MSC-derived PGE2 predominantly protected quiescent HSPCs from IR-induced apoptosis, direct pharmacological elevation of cAMP with Forskolin/IBMX (FSKN/IBMX) effectively shielded both quiescent and cycling HSPCs, significantly enhancing their engraftment and self-renewal. Mechanistically, cAMP pathway activation reduced pro-apoptotic ASPP1 and PUMA expression, elevated p21, and stabilized anti-apoptotic MCL1 and BCL-XL proteins. Collectively, our study uncovers an MSC-driven PGE2/CREB signaling pathway critical for human HSPC regeneration, highlighting pharmacological modulation of this axis as a promising strategy to mitigate DNA damage-induced myelosuppression and improve transplantation outcomes.

## Introduction

Hematopoietic Stem and Progenitor Cells (HSPCs) are rare bone marrow (BM) cells (<0.1% of nucleated cells) positioned at the apex of the hematopoietic hierarchy. They possess the unique ability to self-renew and differentiate into multiple blood cell lineages. These regenerative properties are crucial for lifelong blood production, the success of autologous or allogeneic HSPC transplantation, and the regeneration of hematopoiesis following ionizing radiation (IR) or chemotherapy^1,2^.

The hematopoietic system is among the most radiosensitive tissues in our body. DNA damage to HSPCs critically limits hematopoietic regeneration by inducing apoptosis, impairing self-renewal, and causing aberrant differentiation. Patients with inherited defects in DNA damage response genes, as well as murine models of these diseases, frequently exhibit marked genomic instability, progressive BM failure due to impaired HSC function, and an increased incidence of hematopoietic malignancies^3^. A comprehensive understanding of the molecular pathways that regulate HSPC survival, maintenance, and regeneration under genotoxic stress is critical for developing therapeutic strategies to mitigate these adverse outcomes.

Prior work, including our own, have established that irradiated human HSPCs undergo rapid apoptosis and exhibit a pronounced loss of long-term repopulation capacity^4–6^. Genetic approaches have implicated key regulators such as p53, ASPP1, BCL-2, and MCL1 in the HSPC response to genotoxic injury^4,7–9^. Notably, while permanent p53 inactivation transiently rescued irradiated HSPCs, BCL-2 overexpression preserved self-renewal, highlighting the critical role of p53 in maintaining HSPC genomic integrity for long-term regeneration^4^.

The heightened susceptibility of hematopoietic cells to genotoxic stress has driven pioneering studies in murine models to identify factors that promote HSPC regeneration. These studies implicated numerous BM niche-secreted factors in mitigating DNA damage-induced HSPC injury, including SCF^10,11^, EGF^12,13^, VEGF^14^, Angiopoietin^15^, Pleiotrophin^16–18^, Dickkopf-1^19^, Angiogenin^20^, FGF^21,22^, and PGE2^23^. However, when tested on human HSPCs, key factors such as SCF, FLT3L, TPO, EGF and PGE2 failed to inhibit DNA damage-induced apoptosis *in vitro* or protect human hematopoiesis in immunodeficient xenograft models^24^. These discrepancies underscore species-specific differences in irradiation susceptibility^25^, which can stem from variations in transcriptional responses^26^, cytokine requirements, transformation susceptibility, telomere/telomerase biology, and the inbred nature of murine strains, complicating the direct extrapolation of rodent findings to humans^27–29^.

The BM microenvironment consists of a variety of non-hematopoietic cells that engage in dynamic crosstalk with HSPCs within specialized niche structures^30,31^. Mesenchymal stromal cells (MSCs), a critical niche component, support HSPC long-term regenerative capacity in both *in vivo* and *ex vivo* settings^32–35^. Substantial research has explored MSCs therapeutic potential in protecting the hematopoietic system from IR-induced injury^34–42^. While these studies have demonstrated increased survival of irradiated hematopoietic cells in co-culture, the identity of MSC-derived pro-survival factors, the molecular pathways engaged in human HSPCs, and their functional impact *in vivo* remains largely unknown.

Prostaglandin E2 (PGE2) is a bioactive lipid that regulates vertebrate HSPCs via EP2 (PTGER2) and EP4 (PTGER4) Gs-protein-coupled receptors, activating the cyclic AMP (cAMP) signaling pathway^43–45^. The cAMP-responsive element-binding protein (CREB), a master transcription factor activated by elevated cAMP levels, mediates PGE2-dependent transcriptional responses^46^. Importantly, short-term (2-hour) PGE2 exposure enhanced HSPC migration, survival under cytokine deprivation, homing ability, and human chimerism in xenograft models^47–49^. However, this regimen failed to improve neutrophil recovery in a clinical trial^48^. Likewise, prolonged (24–48 hours) ex vivo PGE2 stimulation before xenotransplantation did not enhance human chimerism^50–52^. Notably, pre-IR injection of PGE2 rescued murine HSPC functionality^53^, whereas the opposite effect was observed in humanized mice^24^. These findings suggest that while PGE2/cAMP signaling axis exerts pleiotropic effects on isolated HSPCs, its role in mediating MSC-driven pro-survival functions and its precise molecular mechanisms in human HSPC regeneration after DNA damage remain unresolved.

The advances in methodologies for isolating, manipulating, and analyzing primary human HSPCs, particularly in co-culture systems with MSCs and integrated transcriptomic approaches, provide a unique opportunity to dissect the molecular mechanisms underpinning HSPC regeneration following DNA damage. Our study demonstrates that MSC-secreted PGE2 activates cAMP signaling via EP2 and EP4 receptors in HSPCs, preventing IR-induced apoptosis by rapidly modulating ASPP1, p21, and PUMA expression. Notably, pharmacological activation of cAMP signaling with small-molecule agonists replicated MSC-mediated protection, strongly enhancing HSPC functionality and self-renewal.

## Results

### MSCs promote stemness and anti-inflammatory transcriptional program in human HSPCs

Human HSPCs are acutely sensitive to ionizing radiation (IR), resulting in apoptosis and loss of hematopoietic reconstitution potential^4,5,54^. To decipher the molecular mechanisms underlying MSC-mediated protection against genotoxic stress, we established a niche mimicking human CD34+-MSC co-culture system (Figure 1A). As expected, upon irradiation the more immature fraction defined here as CD34+CD38-CD45RA-HSPCs showed a significant increase in Annexin-positive apoptotic cells, whereas co-culture with human BM-MSCs completely prevented apoptosis (Figure 1B). Due to the limited availability of primary human BM-MSCs, we used the well-characterized murine OP9M2 MSC cell line to support and extend results obtained with human MSCs^55^. Notably, OP9M2 cells are known to support human hematopoiesis ex vivo^56^ and we independently confirmed their surface profile characteristics to MSCs (Supplemental Figure 1A). Co-culturing CB-derived HSPCs with OP9M2 prevented apoptosis induced by IR or Etoposide treatments (Figure 1B; Supplemental Figure 1B-C).

**Figure 1.**
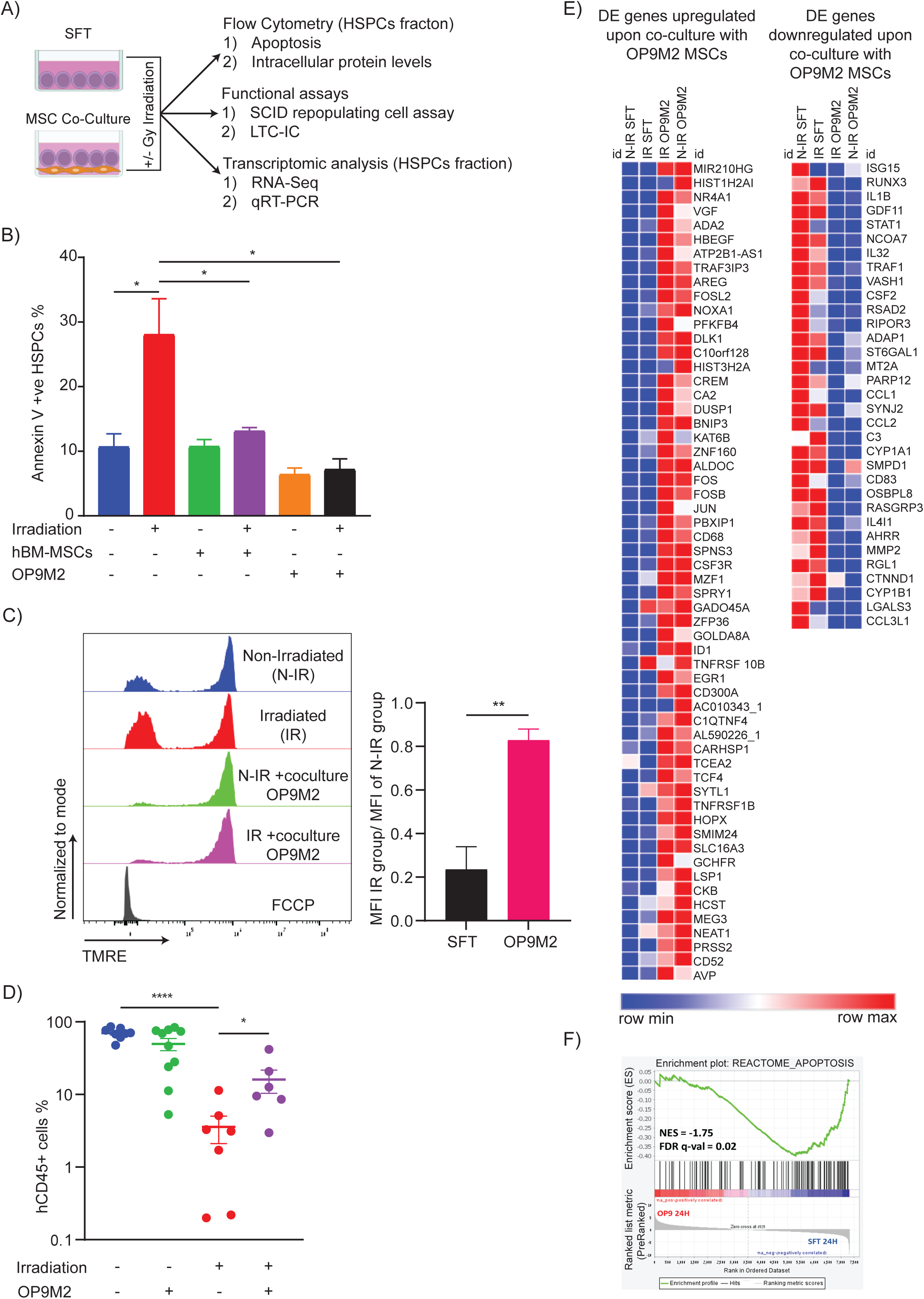
MSCs protect human HSPCs from DNA damage-induced apoptosis. **(A)** Schematic of the experimental design to evaluate HSPC survival and functionality following co-culture with MSCs post-irradiation. **(B)** Flow cytometric analysis of apoptosis in HSPCs cultured for 24 hours—either in cytokine-supplemented medium alone or on human BM-derived MSCs or murine OP9M2 cells—following 3 Gy irradiation, as assessed by Annexin V staining (n=3). **(C)** Analysis of mitochondrial membrane potential in HSPCs under the indicated conditions. Left: Representative TMRE histograms; right: Quantification expressed as the ratio of mean fluorescence intensity (MFI) between irradiated and non-irradiated cells (n=3). **(D)** In vivo assessment of HSPC functionality. CB CD34⁺ cells cultured for 24 hours under the indicated conditions were transplanted retro-orbitally into NSGW41 mice. Left panel: Percentage of human CD45⁺ cells in murine bone marrow 15 weeks post-transplant; right panel: Lineage distribution of engrafted cells. Each symbol represents an individual recipient; horizontal bars denote the mean (n=2). **(E)** Heatmap displaying differentially expressed genes in HSPCs co-cultured with OP9M2 MSCs compared to cytokine-only controls. **(F)** Gene Set Enrichment Analysis (GSEA) plot illustrating the enrichment of apoptosis pathway genes in HSPCs co-cultured with OP9M2 MSCs versus those cultured in serum-free medium with cytokines.

Given the key role of mitochondria in cell functionality and IR-induced apoptosis, we assessed mitochondrial mass (by MitoTracker green) and integrity (by TMRE) in human HSPCs cultured with cytokines or in contact with OP9M2. In the frame of 24 hours, we detected no changes in mitochondrial mass or mitochondrial membrane potential between HSPCs cultured alone or in presence of MSCs. In contrast, while IR did not affect mitochondrial mass, it induced mitochondrial depolarization in HSPCs cultured alone, correlating with increased apoptosis. When co-irradiated, MSCs preserved mitochondrial polarization in HSPCs, consistent with reduced rate of apoptosis (Figure 1C; Supplemental Figure 1D, 1E). Mitochondrial transfer between MSCs and leukemic cells was implicated as MSC cytoprotective mechanism upon injury^57^. We confirmed that there was no transfer of mitochondria from MSCs to HSPCs, excluding this possibility from MSC mediated protection (Supplemental Figure 2A-C).

To quantitate the impact of MSCs on the functionality of irradiated HSPCs we employed long term culture initiating cell (LTC-IC) analysis *in vitro* and SCID-repopulating cell (SRC) assay *in vivo.* We observed a massive drop in LTC-IC frequency when HSPCs were irradiated alone, which was improved by 5-fold upon similar irradiation exposure of HSPCs-MSCs co-cultures (Supplemental Figure 3A). Similarly, IR of CD34+ cells in monoculture impaired SCID-repopulating ability (mean human chimerism in the bone marrow after 12 weeks: 70% NT vs. 3% IR), while irradiation of the MSC-HSPCs co-culture significantly improved HSPC functionality (mean human chimerism: 50% NT vs. 16% IR) (Figure 1D). Radiation dose (3Gy) or presence of MSCs had no impact on HSPCs multilineage differentiation potential or migration to spleen (Supplemental Figure 2B-D). Overall, these findings demonstrate that BM-MSCs protect human HSPCs from genotoxic stress by preventing apoptosis and preserving their functionality.

To delineate the molecular programs driving the enhanced survival and functionality of human HSPCs upon co-culture with MSCs, we performed genome-wide RNA sequencing of human HSPCs sorted after mono- or co-culture with OP9M2 MSCs. Differential gene expression analysis revealed upregulation of AP-1 transcription factors (FOSL2, FOS, FOSB, JUN) and metabolic regulators (ALDOC, PFKFB4), alongside downregulation of pro-inflammatory genes (IL1B, STAT1, CCL1, CCL2) in HSPCs co-cultured with MSCs (Figure 1E).

Gene Set Enrichment Analysis (GSEA) showed that HSPCs co-cultured with MSCs exhibited downregulation of stress-related gene sets (“Apoptosis,” “Unfolded Protein Response”), proliferation-associated pathways (“Myc Targets V1,” “Oxidative Phosphorylation”), and pro-inflammatory responses (“Interferon Gamma Response,” “Interferon Alpha Response”) (Figure 1F, 2A). Conversely, gene sets linked to primitive HSC subsets, including “Hypoxia,” “Glycolysis,” and LT-HSC stemness-associated signatures^58–60^, were enriched in HSPC-MSC co-cultures (Figure 2A). Intriguingly, GSEA analysis also highlighted that HSPCs isolated from MSCs co-culture exhibit enrichment of the transcriptional programs that characterize highly regenerative EPCR+HSPCs^60^ (Figure 2B). To functionally validate these findings, we assessed EPCR expression, a key anti-inflammatory marker of human HSPCs^60^. Consistent with our transcriptomic data, EPCR expression was significantly upregulated in HSPCs cultured with MSCs compared to those in cytokine only conditions (Figure 2C). Overall, our transcriptomic analysis highlights a strong enrichment of stemness-associated gene sets, expansion of regenerative subpopulations, and increased resilience to DNA damage in HSPCs co-cultured with MSCs.

**Figure 2.**
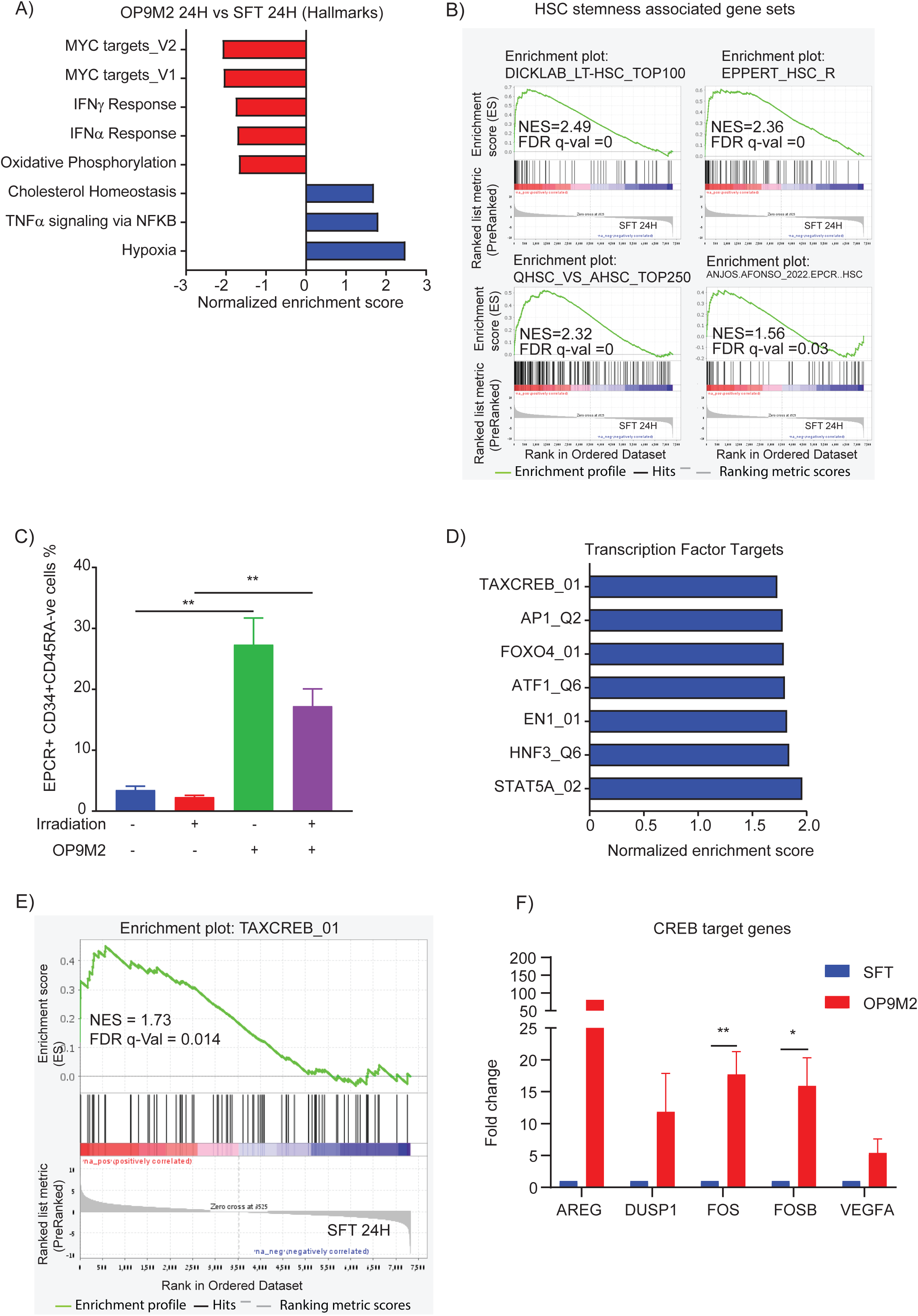
HSPCs co-cultured with MSCs acquire a stemness and anti-inflammatory gene signature. **(A)** GSEA showing significant enrichment of hallmark gene sets in HSPCs co-cultured with OP9M2 MSCs compared to cytokine-only culture. **(B)** Enrichment plots for previously published HSC stemness-associated gene sets **(C)** Flow cytometric quantification of EPCR expression on CD34⁺CD45RA⁻ HSPCs cultured under the indicated conditions for up to 96 hours (n=3). **(D)** GSEA demonstrating significant enrichment of transcription factor target gene sets in HSPCs co-cultured with OP9M2 MSCs relative to cytokine-only conditions. **(E)** GSEA enrichment plot for TAX-CREB target genes in HSPCs co-cultured with OP9M2 MSCs versus cytokine-only culture. **(F)** qRT-PCR analysis validating upregulation of CREB target genes in sorted HSPCs co-cultured with OP9M2 MSCs compared to controls (n=3).

### MSCs activate cAMP/ CREB signaling pathway to protect human HSPCs

To identify upstream transcription factors (TFs) driving gene expression changes in HSPCs upon interaction with MSCs, we performed GSEA using the Transcription Factor Targets (TFT) gene set collection. This analysis revealed MSC-induced activation of several TFs in HSPCs, including STAT5A, HNF3, FOXO4, and notably ATF1, AP-1, and CREB1 all of them belong to same bZIP TF family (Figure 2D, 2E). Among these, CREB1 (hereafter CREB) stood out, as it has been shown in regulating immediate early gene induction and promoting the survival of human CD34+ cells^61^. We therefore hypothesized that activation of the CREB signaling axis might orchestrate the transcriptional changes and pro-survival phenotype observed in HSPCs upon MSC contact.

To test this hypothesis, we first validated our RNA-seq findings via qRT-PCR, confirming strong upregulation of multiple CREB target genes in sorted HSPCs after co-culture with MSCs for 24 hours (Figure 2F; Supplemental Figure 4A). Although multiple signaling pathways can affect CREB transcriptional activity, it is particularly sensitive to the cAMP dependent effectors, e.g. PKA that can activate CREB TF by phosphorylation^62^. To test the possible contribution of cAMP in MSC mediated protection of irradiated HSPCs, we inhibited cAMP-dependent effectors using Rp-8-Br-cAMPs, a competitive cAMP antagonist that prevents CREB phosphorylation^63^. In the presence of Rp-8-Br-cAMPs, the effectiveness of MSCs to protect HSPCs from IR-induced apoptosis was significantly reduced, demonstrating that cAMP dependent signaling required for MSC-mediated radioprotection (Figure 3A).

**Figure 3.**
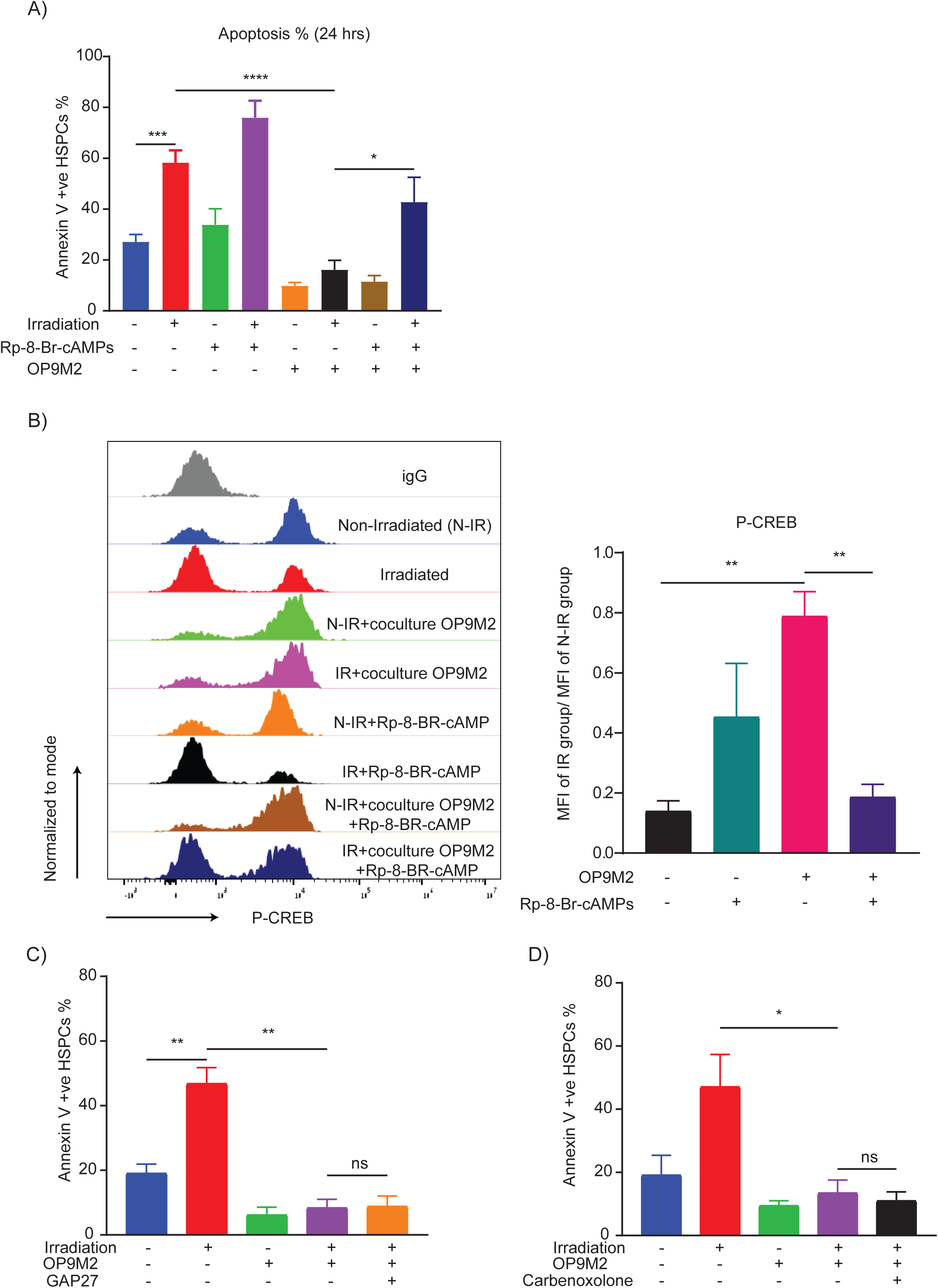
MSCs activate the cAMP/CREB signaling pathway to protect human HSPCs. **(A)** Annexin V-based apoptosis assay showing that CB CD34⁺ cells cultured for 24 hours with MSCs exhibit significantly reduced IR-induced apoptosis compared to cytokine-only controls (n=6). **(B)** Flow cytometric analysis of phospho-CREB (Ser133) levels in HSPCs after 24 hours under the indicated conditions (n=3). **(C–D)** HSPCs cells cultured for 24 hours in the presence of gap junction inhibitors (Carbenoxolone and GAP27) maintain MSC-mediated protection from apoptosis, indicating a gap junction-independent mechanism (n=3).

To determine whether CREB is directly activated upon MSC co-culture, we assessed intracellular levels of phospho-CREB (P-CREB S133). As predicted, P-CREB (S133) levels were elevated in MSC-co-cultured HSPCs, supporting our bioinformatics analysis. Notably, irradiation suppressed P-CREB in HSPCs cultured in SFT, whereas P-CREB levels remained unchanged in the MSC co-culture group (Figure 3B; Supplemental Figure 4B). Importantly, Rp-8-Br-cAMPs abolished P-CREB induction in irradiated HSPCs even in MSC co-culture, reinforcing that MSCs activate CREB in a cAMP-dependent manner (Figure 3B). Together, these findings establish cAMP/CREB signaling as a critical driver of the MSC-mediated pro-survival phenotype in human HSPCs.

Given prior reports of gap junction-mediated cAMP transfer between adjacent cells^64^, we next investigated whether MSCs transmit cAMP to HSPCs through gap junctions. Pharmacological inhibition of gap junctions using Carbenoxolone and GAP27 failed to abrogate MSC-mediated radioprotection, indicating that MSCs enhance HSPC survival through a gap junction-independent mechanism (Figure 3C, 3D).

### PGE2 secreted by BM-MSCs promote HSPCs survival via EP2 and EP4 receptors

To decipher the molecular mechanism by which MSCs block IR-induced apoptosis in HSPCs in a cAMP signaling dependent manner we considered a paracrine stimulation of Gs coupled receptors on the surface of HSPCs. Based on the existing literature we compiled a list of ligands produced by BM MSCs that are capable of stimulating Gs coupled receptors. Next, we examined the expression of their cognate receptors in human HSPCs in our RNA-seq data (Table 1). This analysis pointed out that human HSPCs expressed high level of PGE2 receptors and negligible or undetectable levels of receptors for the additional Gs stimulatory ligands considered (such as Adenosine and PTHrP).

**Table 1.** Potential cAMP activating ligands and their receptors in the context of MSC-HSPC crosstalk. Only ligands capable of stimulating Gs coupled receptors and produced by MSCs were considered.

| BM-MSC-secreted molecules reported to activate Gs coupled GPCRs | Cognate Gs coupled receptors | Receptor expression on CD34+CD38-CD45RA- HSPCs |
| --- | --- | --- |
| Prostaglandin E2 (PGE<sub>2</sub>)<sup>1</sup> | PTGER2 | Low |
|  | PTGER4 | Yes |
| Adenosine<sup>2</sup> | ADORA2A | Not expressed |
|  | ADORA2B | Low |
| Parathyroid hormone-related protein (PTHrP)<sup>3</sup> | PTH1R | Not expressed |

Prostaglandin E2 (PGE2) is a bioactive lipid with pleiotropic effects on hematopoiesis^45,65^. To investigate whether MSC-derived PGE2 contributes to HSPC survival after IR, we measured PGE2 levels in conditioned medium (CM) samples collected from cultures of CD34+ and various MSCs including OP9M2, MS5, and primary hBM-MSCs. While CD34+ cells produced negligible amounts of PGE2, studied MSC cultures secreted at least 10-fold higher levels. We found comparable secretion of PGE2 by established murine MSC lines and primary human BM-MSCs (Figure 4A).

**Figure 4.**
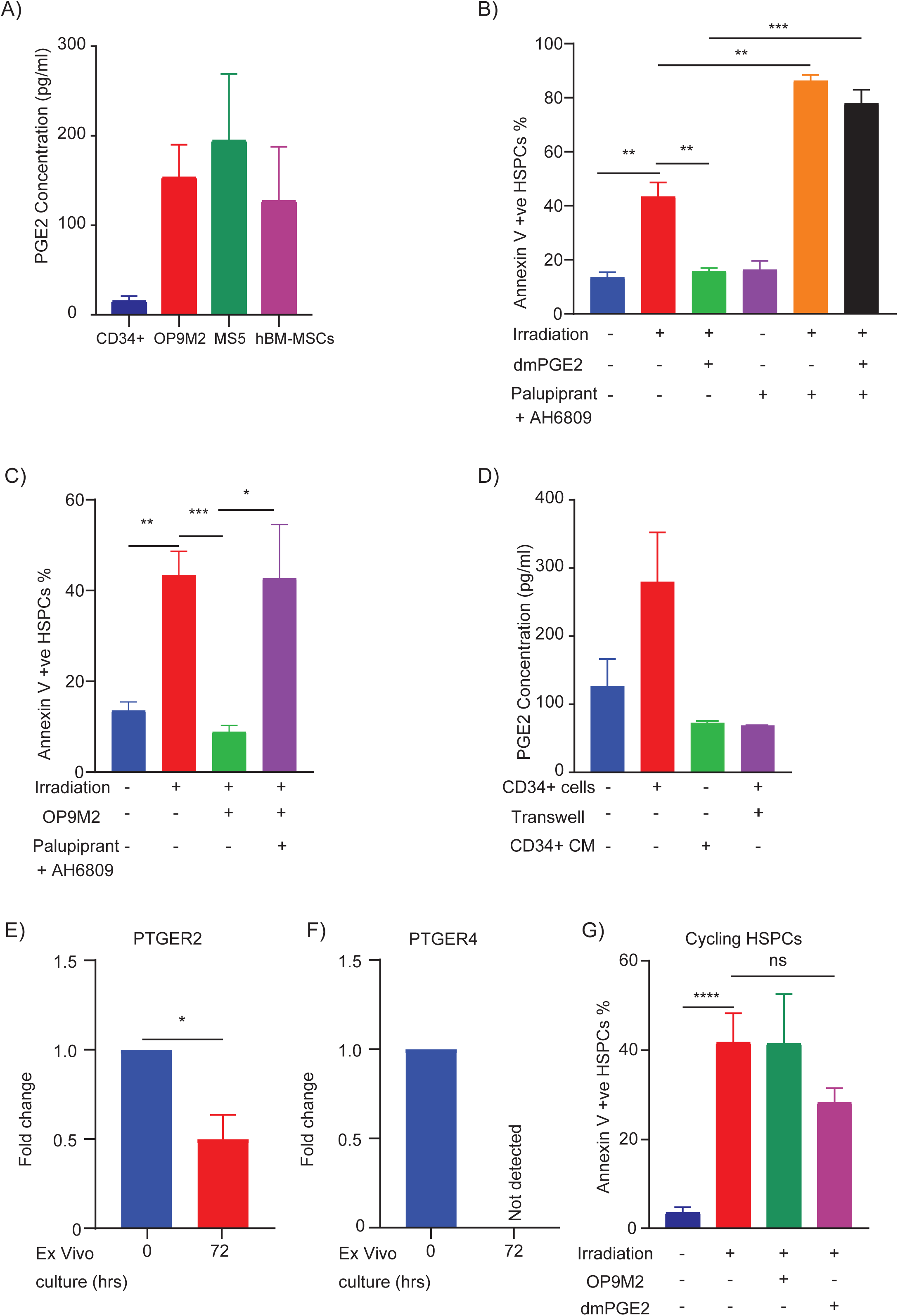
BM-MSCs secrete PGE2 to protect human HSPCs from IR-induced apoptosis. **(A)** ELISA quantification of PGE2 in conditioned medium collected 24 hours after switching OP9M2, MS5, and primary human BM-MSCs from MEMα to SFEM SFT medium (n=4). **(B– C)** Annexin V analysis of apoptosis in HSPCs cultured for 24 hours under the indicated conditions, with or without dmPGE₂ and selective EP receptor antagonists (AH6809 for EP1-3 and Papupiprant for EP4) (n=4). **(D)** Comparison of PGE₂ secretion by OP9M2 MSCs when co-cultured with CD34⁺ cells either in direct contact or separated by a 0.4μm transwell, as measured by ELISA (n=3). **(E–F)** qRT-PCR analysis of prostaglandin E₂ receptor (PTGER1-4) gene expression in freshly isolated versus 72-hour cultured cycling CD34⁺ HSPCs (n=4). **(G)** Annexin V staining of apoptosis in cycling HSPCs (induced by 72-hour culture) following 24-hour exposure to the indicated conditions (n=3).

PGE2 signals through EP2 and EP4 receptors in HSPCs, activating the cAMP pathway^43–45^. To test whether MSC-secreted PGE2 protects HSPCs from IR-induced apoptosis, we used dmPGE2 and selective EP receptor antagonists (EP1-3 antagonist AH6809 and EP4 antagonist Palupiprant) in apoptosis assays. While blocking EP1-3 alone or in combination with EP4 reversed dmPGE2-mediated protection, only their combined inhibition abrogated MSC-mediated protection in both OP9M2 MSCs (Figure 4B, 4C; Supplemental Figure 4C, 4D) and primary hBM-MSCs (Supplemental Figure 4E). To confirm that these EP receptor antagonists did not interfere with PGE2 production by MSCs, we treated OP9M2 MSCs with the inhibitors and measured PGE2 levels in CM. PGE2 production remained unchanged, indicating that the loss of MSC-mediated protection was due to direct inhibition of EP2 and EP4 signaling in HSPCs (Supplemental Figure 4F). Notably, co-culture with CD34⁺ HSPCs enhanced PGE2 secretion by OP9M2 MSCs in a contact-dependent manner, highlighting bidirectional crosstalk between HSPCs and MSCs (Figure 4D).

To further dissect the EP receptor subtype mediating HSPC protection, we assessed expression of *PTGER1-4* genes human HSPCs. Only EP2 and EP4 were expressed in quiescent cord blood HSPCs, consistent with findings by Wang et al.²⁶. Interestingly we found that ex vivo culture of HSPCs led to a strong alteration in PGE2 receptors expression, while PTGER2 (EP2) underwent a 2-fold downregulation, no expression of PTGER4 (EP4) can be detected after 72 hours in culture (Figure 4E, 4F). To substantiate the functional consequences of this receptor downregulation, we tested whether dmPGE2 or OP9M2 continue to block IR-induced apoptosis in cycling HSPCs. In agreement with EP2 and EP4 receptor downregulation on cycling HSPCs, we found loss of pro-survival effect of dmPGE2 treatment prior to IR or co-irradiation on OP9M2 monolayers (Figure 4G). Collectively, these findings establish MSC-derived PGE2 as a key pro-survival factor that protects quiescent, but not cycling HSPCs via EP2 and EP4 mediated receptor signaling, in our ex vivo HSPC-MSC niche model.

### Pharmacological activation of cAMP/CREB signaling pathway protects human HSPCs from IR induced apoptosis

To dissect the molecular role of cAMP signaling in human HSPC response to DNA damage, we employed a pharmacological strategy to elevate intracellular cAMP levels independent of MSC-derived ligands and variable surface receptor expression. Since cAMP levels are tightly regulated by Gs-coupled adenylyl cyclases and phosphodiesterases, we used a combination of Forskolin—a direct and reversible activator of adenylyl cyclases—and IBMX, a broad phosphodiesterase inhibitor to induce a robust and sustained activation of the cAMP/CREB pathway. As expected, Forskolin/IBMX treatment increased P-CREB levels and prevented IR-induced P-CREB loss in HSPCs (Figure 5A, Supplemental Figure 5A).

**Figure 5.**
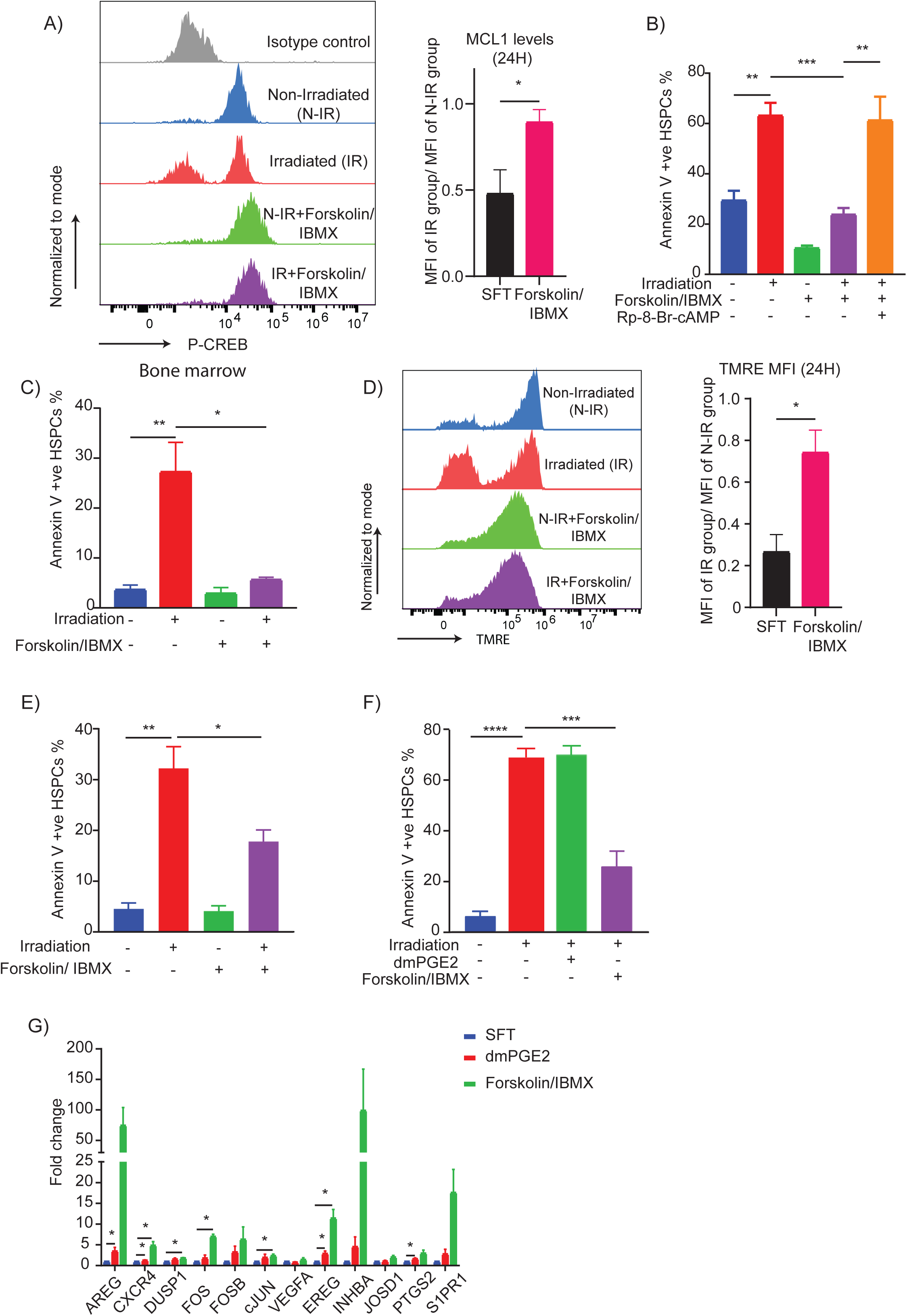
Pharmacological activation of the cAMP/CREB signaling pathway protects human HSPCs from IR-induced apoptosis. **(A)** Flow cytometric quantification of phospho-CREB (Ser133) levels in HSPCs after 24 hours of treatment with Forskolin/IBMX, demonstrating sustained CREB activation post-irradiation (n=4). **(B)** Annexin V staining showing that Forskolin/IBMX pre-treatment significantly reduces IR-induced apoptosis in HSPCs; this protective effect is abrogated by the cAMP antagonist Rp-8-Br-cAMPs (n=4). **(C)** Apoptosis analysis in human BM-derived CD34⁺ cells cultured under the indicated conditions for 24 hours (n=3). **(D)** TMRE staining demonstrating preserved mitochondrial membrane potential in Forskolin/IBMX-treated cells: representative histograms (left) and corresponding MFI quantification (right) (n=3). **(E)** Annexin V analysis of apoptosis in cycling HSPCs (pre-cultured for 72 hours) after 24-hour exposure to the indicated treatments (n=3). **(F)** Quantification of apoptosis in HSPCs cells cultured for 72 hours under the indicated conditions (n=3). **(G)** qRT-PCR analysis of CREB target gene expression in CD34⁺ cells treated with either dmPGE2 or Forskolin/IBMX for 20 hours (n=4).

We next examined whether Forskolin/IBMX-mediated CREB activation could protect HSPCs from IR-induced apoptosis. Remarkably, pre-treatment with Forskolin/IBMX conferred robust radio- and chemo-protection in both neonatal and adult HSPCs (Figures 5B, 5C; Supplemental Figures 5B, 5C). This effect was abolished by the cAMP antagonist Rp-8-Br-cAMPs, confirming the requirement of cAMP-binding proteins in mediating HSPC survival (Figure 5B). Furthermore, Forskolin/IBMX prevented mitochondrial depolarization and in irradiated HSPCs, mimicking the protective effects of MSCs (Figure 5D)

Since Forskolin/IBMX activates CREB signaling independently of EP2/EP4 receptors, we tested its efficacy in protecting cycling HSPCs. Unlike dmPGE2, which provided only transient protection, Forskolin/IBMX protected both cycling and quiescent HSPCs for up to 72 hours post-irradiation (Figures 5E, 5F).

To delineate differences in cAMP/CREB target gene activation between dmPGE2 and Forskolin/IBMX, we performed a time-course analysis. Both treatments induced comparable CREB target gene expression at early time point (3 hours) (Supplemental Figure 5E). However, Forskolin/IBMX elicited a more sustained transcriptional response at later time points (20 hours), suggesting prolonged activation of the cAMP/CREB pathway (Figure 5G). Collectively, these findings establish the cAMP/CREB axis as a critical regulator of DNA damage-induced apoptosis in human HSPCs and highlight Forskolin/IBMX as a potent pharmacological strategy to enhance HSPC resilience to genotoxic stress.

### Pharmacological activation of cAMP/CREB signaling pathway enhances self-renewal ability of human HSPCs

To evaluate the physiological impact of pharmacologically elevating cAMP levels in human HSPCs, we transplanted control- and Forskolin/IBMX-treated HSPCs into NSGW41 mice^66^ (Figure 6A). Forskolin/IBMX pre-treatment significantly enhanced human engraftment in murine bone marrow compared to cytokine-only treated cells (Figure 6B; Supplemental Figures 6A, 6B). Notably, 3Gy irradiated CD34+ cells cultured with cytokines alone failed to establish hematopoietic grafts in most recipients (4 out of 11), with a mean engraftment of 0.5%. In contrast, Forskolin/IBMX pre-treatment enabled successful engraftment in 9 out of 15 recipients, with a mean engraftment of 3.3%—a 6.6-fold increase over the irradiated, cytokine-treated group (Figure 6B). The improved human chimerism in

**Figure 6.**
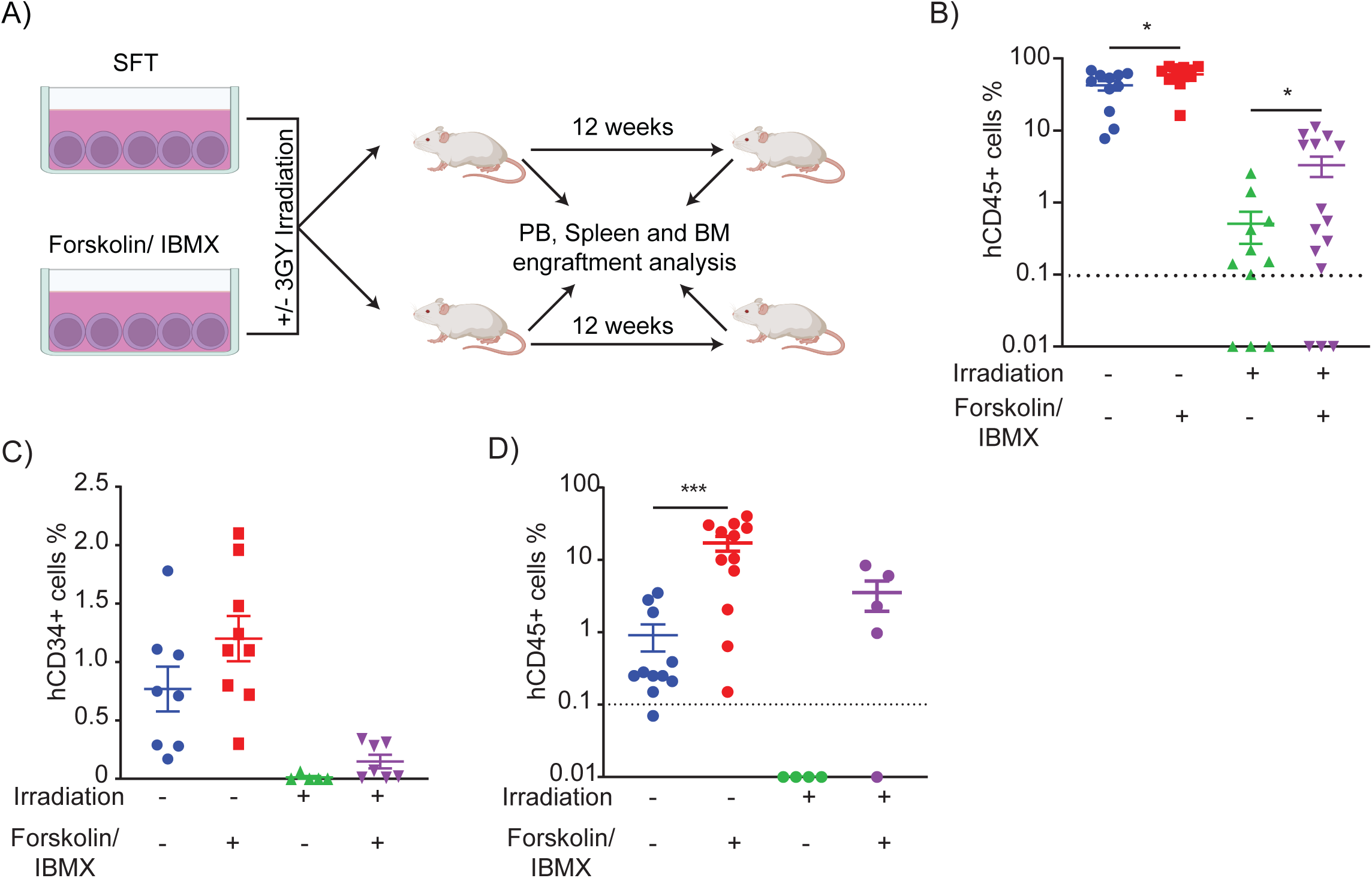
Pharmacological activation of the cAMP/CREB signaling pathway enhances engraftment and self-renewal of human HSPCs. **(A)** Schematic representation of the experimental design to assess HSPC functionality following Forskolin/IBMX treatment. **(B)** Human chimerism in the bone marrow of NSGW41 mice 15 weeks after retro-orbital injection (50,000 CD34+ cells/ mouse) of control or Forskolin/IBMX-treated hCD34⁺ cells (each symbol represents an individual recipient; horizontal bars denote the mean; n=11–15). **(C)** Quantification of immature (hCD34⁺) cell frequency in the bone marrow of mice transplanted with control versus Forskolin/IBMX-treated CD34⁺ cells (n=2). **(D)** Human chimerism in the bone marrow of secondary recipient mice following transplantation of total bone marrow from primary recipients, demonstrating enhanced multilineage regeneration by Forskolin/IBMX-treated HSPCs (each symbol represents an individual recipient; horizontal bars denote the mean).

Forskolin/IBMX-treated HSPCs was accompanied by a modest increase in the proportion of Lin-CD34+ human cells at 12 weeks post-transplantation (Figure 6C). dmPGE2 treatment of CD34+ cells for 24 hours prior to transplantation yielded levels of human chimerism alike the cytokine only group and in agreement with previous reports that used same concentration and exposure length range^67^. dmPGE2 treated and irradiated CD34+ cells demonstrated similarly reduced engraftment abilities comparable to the solvent treated and irradiated group (Supplementary Figure 6C). This comparative *in vivo* analysis underscores previously unknown differences between various cAMP elevating treatments on the enhancement of human HSPC engraftment and regeneration after DNA damage.

To assess the impact of Forskolin/IBMX treatment on HSPC self-renewal, we performed secondary transplantation using bone marrow from engrafted mice. While cytokine-treated HSPCs generated minimal human chimerism (mean engraftment: 0.91%), Forskolin/IBMX-exposed CD34⁺ cells demonstrated robust multilineage regeneration in secondary recipients (mean engraftment: 17.15%) (Figure 6D). Furthermore, irradiated Forskolin/IBMX-treated HSPCs successfully re-established human hematopoiesis in both bone marrow and spleens of secondary recipients (Figure 6D, Supplemental Figure 6D).

To quantitatively assess the frequency of SCID-repopulating cells (SRCs) in cytokine- and Forskolin/IBMX-treated CB CD34⁺ cells, we performed a limiting dilution assay. Poisson distribution analysis revealed an SRC frequency of 1 in 1332 for cytokine-treated cells, compared to 1 in 310 for Forskolin/IBMX-treated cells (Table 2).

**Table 2.** Limiting dilution analysis with Poisson distribution estimates of SCID-repopulating cell (SRC) frequencies in NSGW41 mice, showing an SRC frequency of 1 in 1332 for control and 1 in 310 for Forskolin/IBMX-treated CD34⁺ cells.

| Culture conditions | CD34+ve cells transplanted | Number of mice with >1% human cell chimerism(BM & Spleen) / total number of mice | SRC frequency | 95% confidence interval |
| --- | --- | --- | --- | --- |
| SFT | 500 | 2/5 | 1/1332 | 1/3279 to 1/541 |
|  | 2500 | 4/5 |  |  |
|  | 50000 | 11/11 |  |  |
| Forskolin/ IBMX | 500 | 4/5 | 1/310 | 1/905 to 1/106 |
|  | 2500 | 5/5 |  |  |
|  | 50000 | 12/12 |  |  |

Elevated CXCR4 expression on HSPCs and their increased migration toward SDF-1 following short-term dmPGE2 treatment (2 hours) have been proposed as key contributors to enhanced engraftment potential^44,67^. However, under our experimental conditions—ex vivo exposure to dmPGE2 or Forskolin/IBMX for 24 hours—we observed no improvement in migration toward SDF-1 (Supplemental figure 6E). This finding suggests that enhanced homing is unlikely to be the primary mechanism driving improved engraftment in our system.

Together, these in vivo and in vitro findings establish the cAMP/CREB signaling pathway as a key regulator of human HSPC survival and self-renewal under both homeostatic and genotoxic stress conditions. Moreover, they highlight the potential of specific cAMP-elevating treatments to enhance human HSPC regeneration *in vivo*.

### Pro-survival effects of cAMP agonists are dependent on functional anti-apoptotic MCL1/BCL-XL proteins and altered p53/ASPP1 program

Alterations in p53 transcriptional targets and the balance between pro- and anti-apoptotic BCL-2 family proteins have been implicated in HSPC death following irradiation^68^, yet the precise dynamics of these proteins during apoptosis in human HSPCs remain poorly defined. To that end, we studied the impact of Forskolin/IBMX on the expression of p53 dependent apoptosis regulators and its transcriptional targets in HSPCs. We reveal that cAMP agonist treatment modulated key regulators of the p53 pathway. At 3 hours post-treatment, we observed downregulation of ASPP1, MDM2 and PUMA, along with increased p21 expression (Figure 7A). These data indicates that cAMP-elevating agents rapidly reprogram p53 transcriptional response via downregulating ASPP1 levels.

**Figure 7.**
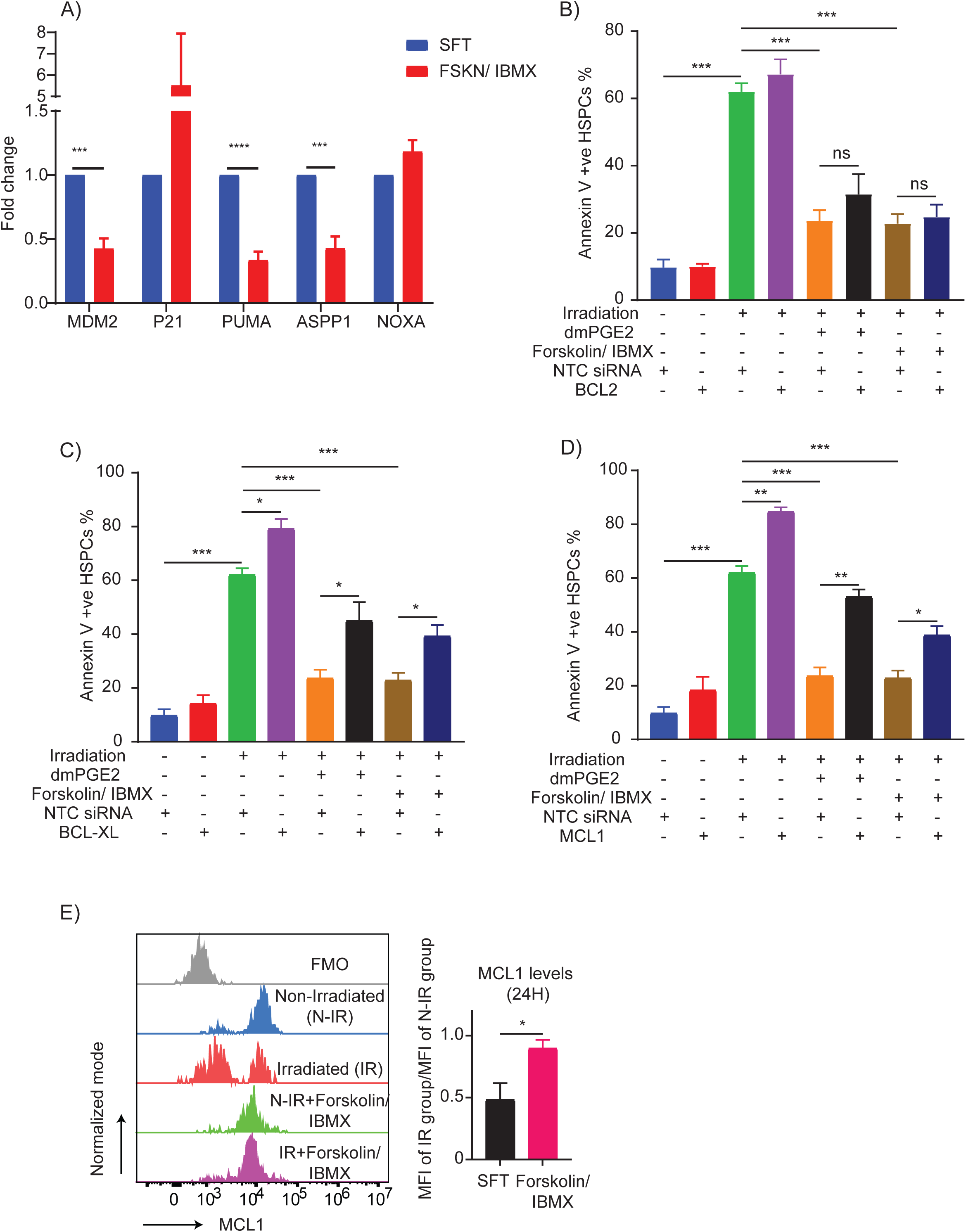
Pro-survival effects of cAMP/CREB signaling depend on functional MCL1 and BCL-XL proteins. **(A)** qRT-PCR analysis of P53 target gene expression in CD34⁺ cells treated with Forskolin/IBMX for 2 hours (n=4). **(B–D)** CB CD34⁺ cells were nucleofected with non-targeting control (NTC) or siRNA pools targeting BCL2 (A), BCL-XL (B), or MCL1 (C). After 24 hours in SFEM with cytokines, cells were exposed to the indicated conditions for an additional 24 hours. Apoptosis in HSPCs was quantified by Annexin V staining (n=3). **(E)** Flow cytometric analysis of MCL1 protein levels in CD34⁺ cells cultured with or without Forskolin/IBMX for 24 hours. Left: Representative histograms; right: Quantitative MCL1 expression (n=6).

To determine whether the radioprotective effects of PGE2 and Forskolin/IBMX in human HSPCs are mediated via specific pro-survival BCL-2 family members, we assessed expression at several time points after treatment. At 3 hours, qRT-PCR analysis demonstrated that both dmPGE2 and Forskolin/IBMX reduced transcript levels of BCL2 and BCL-XL while markedly upregulating MCL1 (Supplemental Figures 7A–7C). By 20 hours, the downregulation of BCL2 persisted; however, BCL-XL and MCL1 transcript levels returned to those observed in untreated cells indicating a rapid but transient transcriptional shift favoring early MCL1 induction. (Supplemental Figures 7D–7F).

To assess the functional relevance of these transcriptional changes, we performed siRNA-mediated knockdowns of BCL2, BCL-XL, and MCL1 prior to irradiation. Notably, while dmPGE2 and Forskolin/IBMX treatments maintained robust radioprotection in HSPCs with BCL2 knockdown, protection was lost upon silencing either MCL1 or BCL-XL (Figures 7B– 7D). These findings indicate that, despite an early downregulation of BCL-XL transcripts, the survival benefit conferred by these treatments is critically dependent on both MCL1 and BCL-XL, with BCL2 playing a less pivotal role.

Given that mRNA levels do not always correlate with protein abundance—especially for proteins like MCL1, which undergo rapid turnover^69^—we next evaluated protein expression at 24 hours post-IR. In irradiated HSPCs, MCL1 protein levels were markedly reduced, likely contributing to radiation-induced apoptosis. Remarkably, treatment with either Forskolin/IBMX or co-culture on OP9M2 MSCs effectively preserved MCL1 protein levels (Figures 7E, Supplemental Figure 8C, 9A). In contrast, BCL2 and BCL-XL protein levels remained relatively unchanged (Supplemental Figures 8A, 8B, 9A–9C), suggesting that selective mechanisms are engaged to stabilize MCL1 under these conditions.

Survivin (BIRC5), a known inhibitor of caspase-dependent apoptosis, has been previously shown to be induced in PGE2 pulse treated human CD34+ cells^67^. We found no changes in Survivin mRNA or protein expression in CD34+CD38-CD45RA-HSPCs upon dmPGE2, FSKN/IBMX or MSCs. However, we do find Survivin expression decreased in IR-exposed HSPCs but remained stable in irradiated HSPCs co-cultured with OP9M2 MSCs (Supplemental Figures 10A, 10B) and FSKN/IBMX treatment (Supplemental Figures 10C-10D).

In summary, our integrated analysis demonstrates that dmPGE2 and Forskolin/IBMX conferred radioprotection depends on the stabilization of MCL1 and BCL-XL proteins in human HSPCs. This dual requirement underscores the complexity of IR induced apoptosis pathway regulation and also highlights the importance of a coordinated anti-apoptotic response by cAMP elevation in mitigating IR-induced stress.

## Discussion

Our results establish that activation of cAMP signaling pathway potently inhibits DNA damage-induced apoptosis and enhances the long-term regenerative potential and self-renewal of human HSPCs. In an ex vivo niche-mimicking model, MSCs secrete PGE2 upon contact with HSPCs, engaging EP2 and EP4 receptors to suppress IR-induced apoptosis. Pharmacological modeling revealed that both synthetic PGE2 and the Forskolin/IBMX (FSKN/IBMX) combination mitigate early apoptosis, but only FSKN/IBMX restored the repopulation capacity of damaged HSPCs in both primary and secondary transplantation assays. Mechanistically, cAMP agonists downregulated pro-apoptotic ASPP1 and PUMA, and upregulated MCL1 and p21, aligning with reduced apoptosis and enhanced survival. These findings demonstrate that physiological or pharmacological activation of cAMP signaling ameliorates IR-induced functional loss of human HSPCs.

Dissection of the MSCs impact on the transcriptome of irradiated human HSPCs is unique to this study and extends prior knowledge obtained from murine HSPCs isolated after whole body IR^8,53^. Our transcriptomic analyses revealed that MSC contact induces a distinct gene expression program in HSPCs, characterized by upregulation of stemness-associated transcription factors (e.g., FOS, JUN, EGR1 and CREB) and a concurrent suppression of pro-inflammatory, stress-related and proliferative pathways (e.g. IFNa, IFNg and MYC). Abrupt transcriptional induction of multiple stress pathways in human HSPCs ex vivo was recently proposed to negatively impact expansion and functionality^56,70^. On the other hand, our transcriptomic analysis reveals that MSCs can stabilize human HSPCs ex vivo in the state reminiscent of their freshly isolated counterparts. Interestingly, MSC-induced transcriptional signature in HSPCs was not significantly affected by IR. Our transcriptomic results underscore niche signaling complexity and argue that MSCs prime HSPCs for enhanced resilience to IR and other genotoxic insults.

While dissecting the possible mechanisms by which MSCs can elevate cAMP levels in human HSPCs, we ruled out the involvement of mitochondrial transfer or gap junction mediated exchange^57,64^. In contrast, we reported concurrent elevation in the secretion of the physiological cAMP agonist PGE2 by human and murine MSCs contacting HSPCs in blocking IR-induced apoptosis and promoting regeneration. MSCs hematopoietic supportive activity is primarily linked to secretion of molecules such as CXCL12, SCF, ANGPT1 and Osteopontin amongst others. PGE2 and its synthetic analogue dmPGE2 have shown to play a positive role in HSPC survival^43,67^. Our data indicates that PGE2 secretion by MSCs chiefly accounts for their ability to suppress IR and Etoposide induced apoptosis. Indeed, blocking the function of PGE2 receptors on HSPCs and the inability of MSCs to protect EP2/EP4 non-expressing expanded HSPCs underscores the importance of BM-MSC derived PGE2 in regulating HSPC survival. Further understanding of the mechanisms regulating PGE2 production and expression of its receptors on HSPCs will be key for advancing MSC- or PGE2-based therapies aimed to improve HSPCs recovery.

Although the detailed mechanisms that account for the cell type specific control of DNA damage induced apoptosis by cAMP are still elusive, our results reveal that cAMP agonists can repress ASPP1 - the endogenous stimulator of p53-dependent apoptosis in both human and murine HSPCs^4,8^. In agreement with the established role of ASPP1 in modulation of p53 transcriptional response we observed coordinated decline in the expression of pro-apoptotic (PUMA) and upregulation of pro-survival genes (e.g MCL1 and p21) upon HSPC treatment with cAMP agonists. This shift in p53 transcriptional response also provides plausible molecular mechanism for the observed block in IR-induced apoptosis (via decreased PUMA) and early cycling (via elevated p21) of mouse HSPCs upon injection of PGE2^53^. Although cAMP agonists resulted only in partial blocking of PUMA induction, murine models posits that even 50% reduction in PUMA expression suffices to significantly protect from the lethal IR-hematotoxicity^6,71^. Elevated activity of CRE-binding factors (e.g., CREB, ATF1 and AP1 complex) in HSPCs from MSCs co-culture or after treatment with cAMP agonists (this study) or of SIRT1^72^ can additionally contribute to the diminished IR-induced apoptosis in HSPCs. Indeed, efficient transactivation of p53, as well as of CREB and NF-KB target genes, are dependent on their binding to p300 and CBP histone acetyltransferases^73^. Given limited cellular amounts of p300/CBP, higher levels of active CREB, as indicated by Ser133 phosphorylation in HSPCs, might destabilize pro-apoptosis gene regulatory programs^74^, and tilt HSPCs cell fate towards survival.

Our mechanistic studies further underscore the differential importance of anti-apoptotic BCL-2 family proteins in this process. Although early transcriptional responses to cAMP elevated treatments included a transient downregulation of BCL2 and BCL-XL, functional assays demonstrated that the radioprotective benefits of both PGE2 and Forskolin/IBMX depend critically on the stabilization of MCL1 and the activity of BCL-XL. Notably, while BCL-XL protein levels did not decrease immediately following irradiation, it is likely that BCL-XL undergoes rapid deamidation as an immediate response to IR stress^75,76^—a modification that may impair its function—and that its protein levels may decline at later time points. siRNA-mediated knockdowns confirmed that disruption of either MCL1 or BCL-XL, but not BCL-2, abrogated the protective effects of PGE2 and FSKN/IBMX, emphasizing the need for a coordinated anti-apoptotic response in mitigating IR-induced damage. In addition to mediating cAMP pro-survival effects as uncovered here, elevated dependency of HSPCs on both BCL-XL and MCL-1 for their survival after DNA damage, as uncovered here, and by Erlacher group^7,77^, represents a human specific phenomenon, as mouse counterparts dependent only on MCL-1^78^ and would predict severe hematotoxicity upon clinical targeting with selective inhibitors.

Importantly, despite efficient IR-induced apoptosis abrogation in HSPCs by cAMP agonists ex vivo, total human engraftment (hCD45+ cells) still decreased by 50% relative to non-IR control, indicating in vivo reactivation of DNA damage induced mechanisms limiting HSPC proliferation. Remarkably, our re-transplantation experiments demonstrated that even few FSKN/IBMX pre-treated irradiated HSPCs preserved significant self-renewal ability that was lost completely in the cytokine only cultured irradiated HSPCs. Moreover, they chart a critical time window in which modulation of DNA damage induced apoptosis can play a critical role for preservation of human HSPC functionality *in vivo*. These findings also independently support a concept obtained in transgenic animals with switchable p53^79^ or deleted PUMA^80,81^ suggesting that while transient inactivation of p53-dependent apoptosis can rescue myelosuppression, its delayed functional restoration suffices to suppress IR-induced and compensatory proliferation driven carcinogenesis.

While agents like dmPGE2 have shown promise in murine models^43,53,72^, our data indicates that direct modulation of the cAMP/CREB axis—via compounds such as Forskolin/IBMX— may provide a more robust and sustained protective effect in human HSPCs. Indeed, pulse treatment of murine and human HSPCs with dmPGE2 which also works via activating cAMP pathway, showed enhanced engraftment potential in some preclinical and in a phase I clinical trial studies^48^ and was mainly attributed to improved SDF-1 migratory capacity driven by elevated CXCR4 expression on HSPCs. On the other hand, we uncovered that different cAMP stimulating regiment (e.g. FSKN/IBMX) can greatly elevate primary and secondary transplantation potential of human HSPCs despite comparable homing ex vivo and progenitor numbers in vivo. Although precise cellular and molecular mechanisms accountable for these effects on HSPCs remain unclear, more sustained impact on gene expression by FSKN/IBMX in comparison to dmPGE2 suggest that certain cAMP elevating treatments might specifically affect human HSPCs self-renewal abilities and could serve as an effective strategy to enhance HSPC survival and function prior to transplantation.

In conclusion, our integrated molecular and functional analyses establish that activation of cAMP/CREB signaling pathway strongly potentiates human HSPCs regenerative abilities in particular after DNA damage via decreasing ASPP1/p53 transcriptional activity and preserving vital MCL1 and BCL-XL protein levels. We predict that focusing on the pharmacological modulation of ASPP1/p53 axis in primary human HSPCs may be more directly relevant for therapeutic strategies aimed to mitigate DNA damage associated myelosuppression, enhance stem cell expansion, genome editing efficacy and transplantation potential without promoting transformation.

## Materials and Methods

### HSPCs purification

Cord blood and bone marrow units were obtained according to procedures approved by the institutional review boards of the Sheba Medical Centre, Rabin Medical Center, Tel Aviv Sourasky Medical Center and Tel Aviv University. Informed consent was obtained from all subjects. Two-six deidentified cord blood units were pooled and mononuclear cells were isolated by density gradient centrifugation. CD34+ cells were enriched by positive selection with MACS CD34+ ultra-pure kit (Miltenyi Biotech) according to the manufacturer’s instructions. Purified cells were stored in liquid nitrogen and used at later time points.

### HSPCs culture

Thawed CD34+ cells were cultured in StemSpan SFEM II serum-free medium (Stem cell technologies) supplemented with recombinant cytokines (Peprotech Asia): SCF (25ng/ml), FLT3L (25ng/ml), TPO (25ng/ml). To elevate cAMP levels, CD34+ cells were treated with Forskolin (10μM, Alomone labs) and IBMX (100μM, Sigma Aldrich). CD34+ cells were treated with following inhibitors– Etoposide (1µM, Sigma Aldrich) S63845 (0.1µM, Apex Biotech), A1155463 (0.1µM, Selleck Chem), Rp-8-Br-cAMPs (100µM, BioLog), Carbenoxolone (100µM, Sigma Aldrich), GAP27 (100µM, Med Chem Express), 16,16-dmPGE2 (10µM, Cayman Chemical), AH6809 (10µM, Med Chem Express), Palupiprant (1µM, Med Chem Express).

### HSPCs and stroma co-culture

OP9M2 murine stromal cells were cultured in MEM-Alpha medium (Gibco, Life Technologies), supplemented with FBS (20%, Biological Industries), Pen/Strep (1%), L-glutamine (1%). Primary Human Bone Marrow stromal cells (BMSCs) were generated from bone marrow aspirate of healthy volunteers after signing an informed consent. BMSCs were isolated by plastic plate adherence (2 × 10^4^ cells per well in 24-well plates incubated overnight) and expanded for 48 hours. Omi-MS5 cells were generated by transducing MS5 cells with pBabe(puro)-Omi-mCherry plasmid (Addgene plasmid# 48685). MS5 and Omi-MS5 cells were cultured in MEM-Alpha medium (Gibco, Life Technologies), supplemented with FBS (10%, Biological Industries), Pen/Strep (1%), L-glutamine (1%).

For co-culture with CD34+ cells, 3*10^4^ OP9M2 cells or 5*10^4^ MS5 cells were plated in 24-well tissue culture treated plate (Greiner Bio-One) and incubated in the stroma medium (see above) for 24 hours. Following the incubation, medium was aspirated, and CD34+ enriched cells were plated on MSCs in StemSpan SFEM II serum-free medium supplemented with cytokines. Cells were maintained in a humidified incubator at 37 °C and CO2 (5%).

### Annexin V apoptosis assay

To induce apoptosis, CD34+ cells cultured in different conditions were irradiated at 3 Gy using a Biobeam gamma-irradiator (Gamma service) or alternatively allowed to cycle for 72 hours before irradiation. For treatment with Etoposide, CD34+ cells were cultured for 24 hours and then treated with Etoposide (1µM) for an additional 48 hours. CD34+ cells were then washed and incubated for 30 mins with antibodies for surface markers – CD45RA BV605 (1:200), CD38 PE/Cy7 (1:100), and CD34 PE (1:100). CD34+ cells were then washed and stained with Annexin V-AF488 (Invitrogen) and Sytox blue dead cell stain (Invitrogen). Cells were analyzed with Cytoflex flow cytometer (Beckman Coulter).

### Intracellular flow cytometry

CD34+ cells were cultured in different conditions as indicated and stained with antibodies for HSPC surface markers - CD45RA BV605, CD38 PE/Cy7 and CD34 PE. Zombie NIR dye (Biolegend) was used to label dead cells. Cells were then fixed with 1.6% Paraformaldehyde and permeabilized in 90% ethanol followed by labelling with AF488-conjugated intracellular antibodies for BCL-2 (1:100), BCL-XL (1:100), MCL-1 (1:100), P-CREB-Ser133 (1:100) and Survivin (1:200).

### Mitochondrial mass & Mitochondrial Membrane Potential analysis

CD34+ cells cultured in different conditions were treated with Verapamil (50μM, Sigma) for 30 mins. MitoTracker Green FM (400nM, Cell Signaling Technology) or TMRE (200nM, Abcam) were added for additional 30 mins. Cells were washed twice, stained with cell surface markers and analyzed by flow cytometry.

### NOD/SCID repopulating cell assay

All animal experimental protocols were approved by the Institutional Animal Care and Use Committee of Tel-Aviv University, Israel. For primary transplantation, human CD34+ cells cultured in different conditions for 24 hours were injected into NSGW41^66^ mice intravenously into the retrobulbar plexus. After 12 weeks, mice were euthanized and BM engraftment was analyzed by flow cytometry using hCD45, hCD33, hCD19 and hCD3 antibodies.

For secondary transplantation, the whole bone marrow of primary mice was reinjected into NSGW41 mice intravenously and the BM engraftment was analyzed by immunostaining and flow cytometry after additional 12 weeks.

The frequency of human SCID repopulating cells (SRCs) was analyzed by limiting dilution analysis. Briefly, increasing doses of SFT or Forskolin/IBMX treated CD34^+^ cells (500, 2500, or 50,000 cells) were intravenously transplanted into NSGW41 mice. 12 weeks after transplantation, the mice were sacrificed and the percentage of human CD45+ cell chimerism was determined by immunostaining and flow cytometry. The HSC frequency was plotted using ELDA software (bioinf.wehi.edu.au/software/elda/).

### In Vitro Limiting dilution analysis

96 well plates (Greiner Bio-One) were covered with 0.1% gelatin solution (Biological Industries) for 20 minutes, and dried open in a sterile hood for 120 minutes. Then, coated wells were seeded with MS5 stromal cells (7000 cells/ well) in 100μl of H5100 medium (STEM CELL technologies) supplemented with 1μM hydrocortisone (Sigma) + 1% P/S. MS-5 cells were irradiated (20 Gy) after seeding into 96 well plates. Twenty-four hours later 10/50/500 non irradiated CD34^+^ cells or 1,500/7,500/15,000 irradiated (3Gy) CD34^+^ cells per well were plated in 100 μl MyeloCult H5100 medium (STEM CELL technologies). Once a week half medium replacement was done. After 5 weeks the H5100 medium was replaced with Methocult H4434 Classic methylcellulose medium (100µl/ well, STEM CELL technologies). Wells that contained at least one colony after 10-14 days were considered positive.

### RNA sequencing

CD34+ cells were cultured in SFT medium or on MSCs for 24 hours, were stained for HSPCs markers and sorted using BD FACS AriaIII cytometer. Total RNA was extracted using TRIZOL reagent (Invitrogen) and quality tested using a 2100 Bioanalyzer (Agilent technologies). Sequencing Libraries were prepared using MARSeq. Single reads were sequenced on 1 lane(s) of an Illumina NOvaSeq_sp. The output was ∼15 million reads per sample.

### Data analysis

Poly-A/T stretches and Illumina adapters were trimmed from the reads using cutadapt^82^; resulting reads shorter than 30bp were discarded. Reads were mapped to the H. sapiens reference genome GRCh38 using STAR^83^, supplied with gene annotations downloaded from Ensembl (and with EndToEnd option and outFilterMismatchNoverLmax was set to 0.04). Expression levels for each gene were quantified using htseq-count^84^, using the gtf above. Differentially expressed genes were identified using DESeq2^85^ with the betaPrior, cooksCutoff and independent filtering parameters set to False. Raw P values were adjusted for multiple testing using the procedure of Benjamini and Hochberg. The Pipeline was run using snakemake workflow engine^86^.

### Gene Set Enrichment Analysis

Gene Set Enrichment Analysis (GSEA)^87^ was done using the GSEA desktop application (Broad Institute). Genes were ranked by the DESeq2 statistic and pre-ranked GSEA was run using the GSEA hallmarks, Transcription Factor Targets (TFT) and other curated gene HSC gene sets from MSigDB database using standard settings. Heatmaps were created using Morpheus (Broad Institute).

### PGE2 detection by ELISA

Conditioned medium from CD34+, OP9M2, MS5, hBM-MSCs or CD34+ - OP9M2 co-cultures were collected 24 hours after treatment. PGE2 levels in the culture medium were determined by PGE2 Enzyme Immunoassay kit according to the manufacturer’s instructions (Cayman Chemicals).

### qRT-PCR analysis

Total RNA was extracted using the RNeasy Micro kit (Qiagen). cDNA synthesis was performed using the qScript cDNA Synthesis kit (Quanta Bio). Real-time quantitative PCR was performed using PerfeCTa SYBR green supermix reagent (Quanta Bio) and analyzed by QuantStudio 5 Real-time PCR system (ThermoFisher). Relative expression was calculated for each gene using by 2^-ΔΔCT method. *GAPDH* was used for normalization.

### siRNA Nucleofection

Dharmacon ON-TARGETplus siRNA pool oligos for non-targeting control, BCL2, BCL-XL and MCL1 were ordered. For electroporation, human 2.5 × 10^5^ CD34^+^ cells/100 µl were resuspended in nucleofection buffer (P3 primary cell kit, Lonza) and were nucleofected with CD34^+^ program using Amaxa 4D nucleofector system (Lonza). Control, si-BCL2, si-BCL-XL, si-MCL1 were used at a final concentration of 2 μM.

### In vitro homing assay

Cord blood-derived CD34⁺ cells cultured with different cAMP elevating treatment for 24 hours were washed and resuspended in cytokine-free medium.

For migration, 300 µl of SFEM II medium containing 300 ng/ml rhSDF-1α (PeproTech) or medium alone (control) was placed in the lower chamber. Plerixafor (25 µM) was added to designated wells for 30 min. A total of 100 µl cell suspension (2 × 10⁴ cells) was added to the upper chamber. Plates were incubated at 37 °C for 4 h. Migrated cells in the lower chamber were stained with CD34 and CD38 antibodies and quantified by flow cytometry.

### Statistical analysis

Statistical analysis was performed with GraphPad Prism 9, using student’s t-test. Statistical significance is defined as p<0.05. Bars represent Mean ± Standard error of mean (SEM) of independent experiments. *P≤0.05, **P≤0.01, ***P≤0.001, ****P≤0.0001.

### Data and materials availability

All raw sequencing data generated in this study have been submitted to the NCBI Gene Expression Omnibus (GEO; https://www.ncbi.nlm.nih.gov/geo/) under the accession number GSE247652.

## Acknowledgments

The authors thank Dr. H.K.A. Mikkola (University of California, Los Angeles, California, USA) for kindly providing the OP9M2 cell line. This work was partially supported by Israel Science Foundation (ISF 3480/19, ISF 1362/20) and Varda and Boaz Dotan Research Center in Hemato-Oncology grants (to MM). This work was performed in partial fulfillment of the requirements for PhD degrees of Siva Sai Naga Anurag Muddineni, the Dr. Miriam and Sheldon G. Adelson Graduate School of Medicine, Faculty of Medicine, Tel Aviv University, Israel. The authors would like to thank Dr. Irena Shur, Dr. Daria Makarovsky and Dr. Rami Khosravi for their technical assistance.

## Author Contributions

SSNAM and MM conceptualized the study, designed and analyzed experiments. SSNAM, CKE, AZ, DR, DSR and RA performed experiments; SSNAM and EW analyzed RNA-seq data; YR, GH, RC, YS, AN, NS, CW and KB prepared and provided study material for experiments; SSNAM and MM wrote the original manuscript; MM provided financial support.

