## Supplemental figure legends for "Mesenchymal Stromal Cells regulate human Hematopoietic Stem Cell survival, engraftment and regeneration via PGE2/cAMP signaling pathway"

### Supplementary figure legends

**Supplemental Figure 1. Characterization of MSCs and initial HSPC apoptotic responses.**

**(A)** Representative histograms illustrating the perivascular marker expression of OP9M2 MSCs. Unstained (isotype control) cells are shown in grey, whereas antibody-stained cells are depicted in red. **(B)** Human BM-derived $CD34^+$ cells were cultured under the indicated conditions for 24 hours and analysed for apoptosis using Annexin V staining (n = 3). **(C)** CB $CD34^+$ cells were cultured for 24 hours in serum-free medium supplemented with SFT cytokines, then treated with 1 µM Etoposide for 48 hours. Apoptosis was assessed by Annexin V staining (n = 3). **(D)** Quantification of mitochondrial membrane potential in HSPCs, as measured by TMRE staining 24 hours after IR and co-culture with OP9M2 MSCs (MFI values). **(E)** CB $CD34^+$ cells were cultured under the indicated conditions for up to 24 hours and stained with MitoTracker Green FM to assess mitochondrial mass.

**Supplemental Figure 2. Analysis of mitochondrial transfer and apoptosis in HSPCs.**

**(A)** CB $CD34^+$ cells were cultured under the indicated conditions for up to 24 hours and analysed for apoptosis by Annexin V staining (n = 3). **(B)** mCherry fluorescence confirming the genetic labelling of MS5 cell mitochondria. **(C)** Representative FACS plots showing mCherry fluorescence in HSPCs cultured alone or in co-culture with mCherry-labelled MS5 cells.

**Supplemental Figure 3. LTC-IC assay and in vivo hematopoietic reconstitution.**

**(A)** CB-derived $CD34^+$ cells were exposed to 3 Gy IR before being cultured for 24 hours in either cytokine-only or MSC co-culture conditions. Cells were then plated for a limiting dilution LTC-IC assay. After five weeks, the medium was replaced with methylcellulose, and wells were scored for colony formation 10–14 days later. The table summarizes LTC-IC frequencies as calculated by L-CALC software. **(B)** Lineage distribution of human cells in the bone marrow of recipient mice. **(C)** Human $CD45^+$ chimerism in the peripheral blood of NSGW41 mice 15 weeks post-transplant. **(D)** Human $CD45^+$ chimerism in the spleen of NSGW41 mice 15 weeks post-transplant.

**Supplemental Figure 4. CREB activation and apoptosis in HSPCs.**

**(A)** Heatmap displaying the expression of leading-edge CREB1 target genes. **(B)** CB $CD34^+$ cells were cultured under the indicated conditions for up to 24 hours and analysed for intracellular phospho-CREB (Ser133) levels. Bar graphs represent the MFI values of P-CREB in HSPCs at 24 hours. **(C–E)** CB $CD34^+$ cells cultured under the indicated conditions for up to 24 hours were analysed for apoptosis using Annexin V staining (n = 3). **(F)** OP9M2 MSCs were cultured under the indicated conditions for 24 hours, and $PGE_2$ concentrations in the supernatants were measured by ELISA (n = 3).

**Supplemental Figure 5. Effects of Forskolin/IBMX on cAMP/CREB activation and apoptosis.**

**(A)** CB CD34$^+$ cells were cultured under the indicated conditions for up to 24 hours and analysed for intracellular phospho-CREB (Ser133) levels (n = 3–4). **(B)** CB CD34$^+$ cells were cultured for 24 hours under the indicated conditions and apoptosis was quantified by Annexin V staining (n = 4). **(C)** CB CD34$^+$ cells were cultured for 24 hours in serum-free medium with SFT cytokines, then treated with 1 µM Etoposide plus 10 µM Forskolin/100 µM IBMX for 48 hours. Apoptosis was assessed by Annexin V staining (n = 3). **(D)** Quantitative analysis (MFI values) of TMRE staining in HSPCs 24 hours after IR and Forskolin/IBMX treatment. **(E)** qRT-PCR analysis of CREB target genes in CD34$^+$ cells treated with dmPGE$_2$ or Forskolin/IBMX for 3 hours (n = 4).

**Supplemental Figure 6. In vivo hematopoietic reconstitution analysis.**

**(A)** Lineage distribution in the bone marrow of transplanted recipient mice. **(B)** Human chimerism in the spleens of mice transplanted with control or Forskolin/IBMX-treated hCD34$^+$ cells (expressed as % hCD45$^+$ cells). **(C)** Human chimerism in the BM of mice transplanted with control or dmPGE2 treated CD34+ HSPCs **(D)** Human chimerism in the spleens of secondary recipient mice following transplantation of bone marrow from primary recipients treated with control or Forskolin/IBMX-treated hCD34$^+$ cells. **(E)** Migratory rate of human HSPCs treated with different cAMP elevating agents for 24hrs towards SDF1a.

**Supplemental Figure 7. qRT-PCR analysis of pro-survival BCL-2 family gene expression.**

CB CD34$^+$ cells were treated with dmPGE$_2$ or Forskolin/IBMX, and gene expression was quantified by qRT-PCR at different time points: **(A)** BCL2 at 3 hours; **(B)** BCL-XL at 3 hours; **(C)** MCL1 at 3 hours; **(D)** BCL2 at 20 hours; **(E)** BCL-XL at 20 hours; **(F)** MCL1 at 20 hours. **(G)** CB CD34$^+$ cells cultured under the indicated conditions for 24 hours were analysed for apoptosis by Annexin V staining (n = 3).

**Supplemental Figure 8. Flow cytometric quantification of anti-apoptotic proteins post-irradiation.**

CB-derived CD34$^+$ cells were cultured for 24 hours after irradiation in SFEM medium with cytokines or in co-culture with OP9M2 MSCs. Intracellular protein levels in HSPCs were measured by flow cytometry: **(A)** BCL2; **(B)** BCL-XL; **(C)** MCL1. Left panels show representative histograms; right panels depict relative expression changes, expressed as the ratio of MFI between IR and non-IR conditions (n = 4 for BCL2 and BCL-XL; n = 6 for MCL1).

---

**Supplemental Figure 9. Analysis of intracellular anti-apoptotic protein levels following Forskolin/IBMX treatment.**

CB-derived CD34$^+$ cells were cultured for 24 hours after irradiation in SFEM medium with cytokines, with or without Forskolin/IBMX treatment. Intracellular levels of the following proteins were assessed by flow cytometry in HSPCs: **(A)** MCL1; **(B)** BCL2; **(C)** BCL-XL. Representative histograms are shown on the left, and relative expression changes (ratio of MFI between IR and non-IR conditions) are shown on the right.

**Supplemental Figure 10. Survivin expression in HSPCs.**

**(A–B)** CB CD34$^+$ cells were cultured under the indicated conditions for up to 24 hours and subsequently analyzed for intracellular Survivin expression in HSPCs. **(C–D)** CB CD34$^+$ cells were cultured under the indicated conditions for up to 24 hours and subsequently analyzed for intracellular Survivin expression in HSPCs.

| **Antigen** | **Clone** | **Fluorophore** | **Dilution** | **Manufacturer** | **Catalog number** |
|---|---|---|---|---|---|
| CD34 | 581 | FITC | 1-100 | Biolegend | 343504 |
| | 581 | PE | 1-100 | Beckman Coulter | A07776 |
| | 8G12 | APC | 1-100 | BD Bioscience | 345804 |
| CD38 | HB7 | PC7 | 1-100 | Biolegend | 356608 |
| CD45RA | HI100 | BV605 | 1-200 | Biolegend | 304134 |
| CD201 | RCR-401 | PE | 1-100 | Biolegend | 351904 |
| | RCR-401 | APC | 1-100 | Biolegend | 351906 |
| CD33 | P67.6 | PE | 1-100 | Biolegend | 366608 |
| CD19 | SJ25C1 | APC | 1-100 | Biolegend | 363006 |
| Lineage cocktail (CD3/14/16/19/20/56) | | FITC | 1-50 | Biolegend | 348801 |
| CD44 | IM7 | PE | 1-100 | Biolegend | 103023 |
| mSCA1 | D7 | FITC | 1-100 | Biolegend | 108105 |
| mCD140a | APA5 | PE | 1-100 | Biolegend | 135905 |
| mCD106 | 429 | APC | 1-100 | Biolegend | 105717 |
| mCD11b | M1/70 | FITC | 1-100 | Biolegend | 101205 |
| mCD31 | 390 | PC7 | 1-100 | Biolegend | 102417 |
| mCD34 | HM34 | APC | 1-100 | Biolegend | 128611 |
| mCD45 | 30-F11 | BV605 | 1-100 | Biolegend | 103140 |
| CD3 | REA613 | FITC | 1-100 | Miltenyi Biotec | 130-114-138 |
| CD45 | J33 | PC7 | 1-100 | Beckman Coulter | IM3548 |
| IgG1 | MOPC-21 | AF488 | 1-400 | Cell Signaling Technology | 4878S |
| IgG1 | DA1E | AF488 | 1-400/1-800 | Cell Signaling Technology | 2975S |
| BCL2 | 124 | AF488 | 1-100 | Cell Signaling Technology | 59422S |
| BCL-XL | 54H6 | AF488 | 1-100 | Cell Signaling Technology | 2767S |
| MCL1 | D2W9E | AF488 | 1-100 | Cell Signaling Technology | 58326S |
| P-CREB (S133) | 87G3 | AF488 | 1-200 | Cell Signaling Technology | 9187S |
| Survivin | 71G4B7 | AF488 | 1-200 | Cell Signaling Technology | 2810S |
| Annexin V | | AF488 | 1-200 | Invitrogen | A13201 |
| | | APC | 1-200 | Invitrogen | A35110 |

| | | | | | |
|---|---|---|---|---|---|
| Sytox Blue | | | 1-1000 | Invitrogen | S34857 |
| Zombie NiR | | | 1-1000 | Biolegend | 423105 |
| TMRE | | | | Abcam | Ab113852 |
| MitoTracker Green FM | | | | Cell Signaling Technology | 9074 |

### List of primers

| | Forward | Reverse |
|---|---|---|
| GAPDH | TTC GTC ATG GGT GTG AAC CA | CTG TGG TCA TGA GTC CTT CCA |
| PTGER1 | TTG GCG GCT CTC GGA | GCC ACC AAC ACC AGC ATT |
| PTGER2 | AGG AGA GGG GAA AGG GTG TC | AAT CGT GAA AGG CAA GGA GC |
| PTGER3 | CCG CAT CAC GAC CGA GAC | AAT CGT GAA AGG CAA GGA GC |
| PTGER4 | ATT CGT CCG CCT CCT TGA | GCC ACC AGG TGG CCC A |
| AREG | TGA GAT GTC TTC AGG GAG TG | AGC CAG GTA TTT GTG GTT CG |
| DUSP1 | TTC TTC CTC AAA GGA GGA TAC G | GTG GGG TAC TGC AGG AAC TG |
| FOS | CGT CTC CAG TGC CAA CTT CA | GGT CCG GAC TGG TCG AGA T |
| FOSB | TTG CAC CTT ACT TCC CCA AC | AGG AGT CCA CCG AAG ACA GA |
| VEGFA | CCA ATC GAG ACC CTG GTG | CAC ACA GGA TGG CTT GAA GA |
| CXCR4 | CCT ATG CAA GGC AGT CCA TGT | GGT AGC GGT CCA GAC TGA TGA |
| cJUN | TCG ACA TGG AGT CCC AGG A | GGC GAT TCT CTC CAG CTT CC |
| EREG | ATC ATG TAT CCC AGG AGA GTC CAG | GAA TCA CGG TCA AAG CCA CAT AT |
| INHBA | TCA CGT TTG CCG AGT CAG GAA C | TGA CAG GTC ACT GCC TTC CTT G |
| JOSD1 | TCC AGG ACA GCA ATG CCT TCA C | CAT GGT GTT TGG AGA CAA CCT CTG |
| PTGS2 | CCC TTG GGT GTC AAA GGT AA | GCC CTC GCT TAT GAT CTG TC |
| S1PR1 | ACG TAG GCT GTG GGA AGA TGA AG | TGG AAA CTT TGG CCT CAG CGA AG |
| ASPP1 | TTGTCCTCTCATTGCACG | AACTTACCCTCTCAGAGC |
| MDM2 | ATCAGCAGGAATCATCGGAC | CCAGGCTTTCATCAAAGGAA |
| PUMA | CCTGGAGGGTCCTGTACAATCT | GGACACAAGAAGAAAACCTTAATGC |
| NOXA | AGCTGGAAGTCGAGTGTGCT | TCCTGAGCAGAAGAGTTTGGA |
| CDKN1A | CGCGACTGTGATGCGCTAATG | GGAACCTCTCARRCAACCGCC |
| BCL2 | TGT GGA TGA CTG AGT ACC TGA ACC | GGAGAAATCCAGAGGCCGCAT |
| BCL-XL | GGA GAA CGG CGG CTG GGA TA | GGC CAC AGT CAT GCC CGT CA |
| MCL1 | CAT TCC TGA TGC CAC CTT CT | TCG TAA GGA CAA AAC GGG AC |
