## Supplemental Figures for "Mesenchymal Stromal Cells regulate human Hematopoietic Stem Cell survival, engraftment and regeneration via PGE2/cAMP signaling pathway"

Supplemental Figure 1

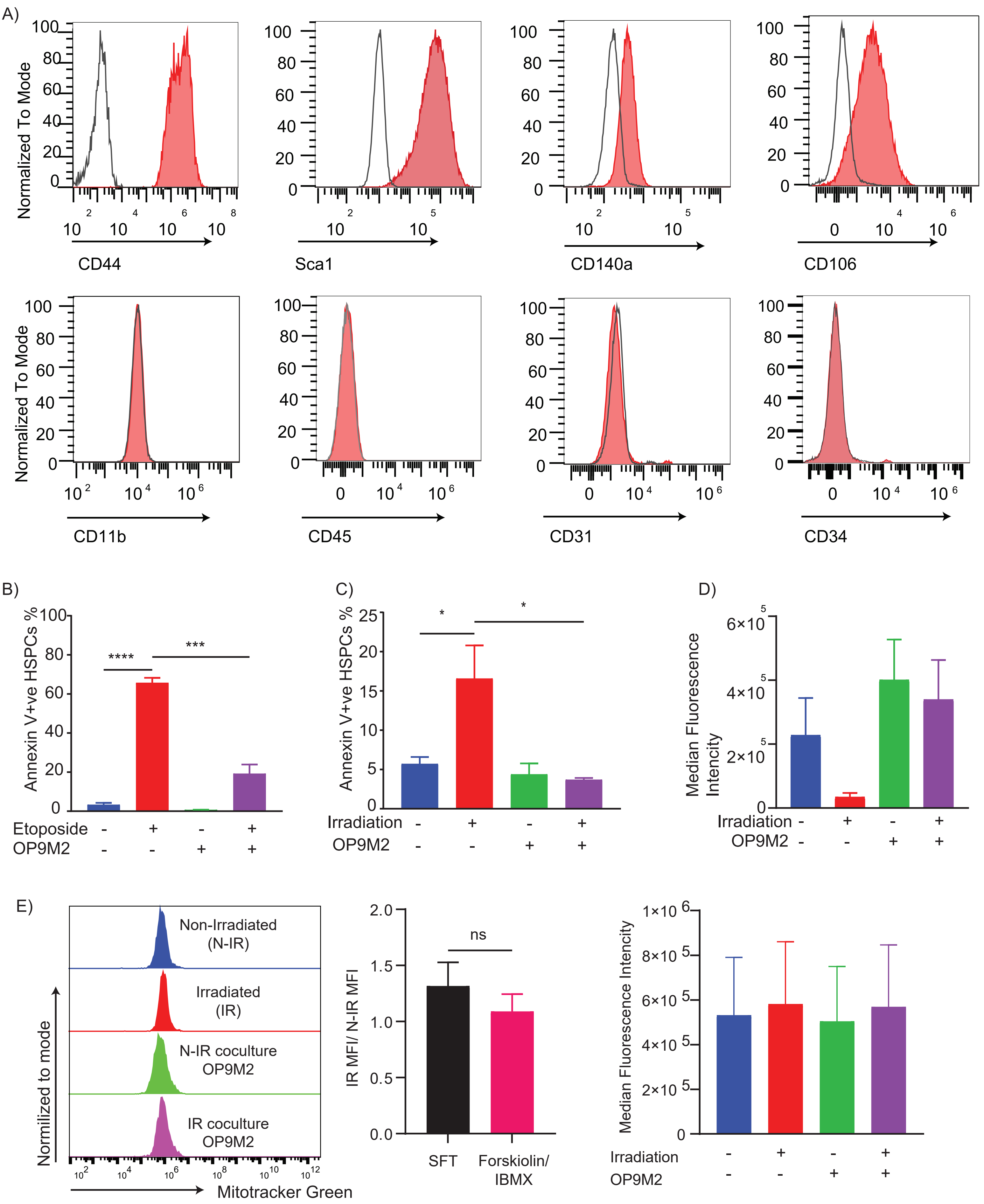

Supplemental Figure 2

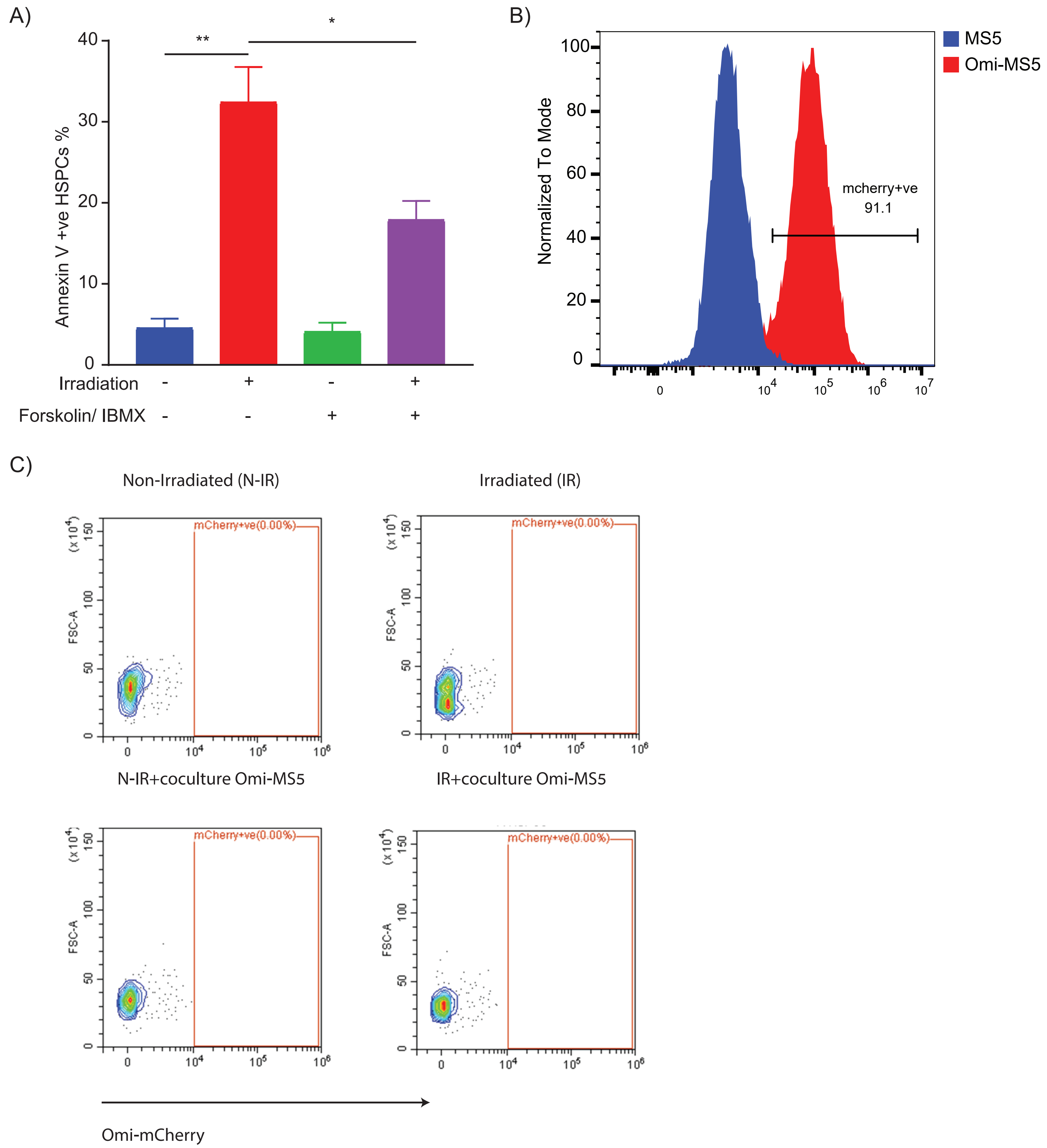

Supplemental Figure 3

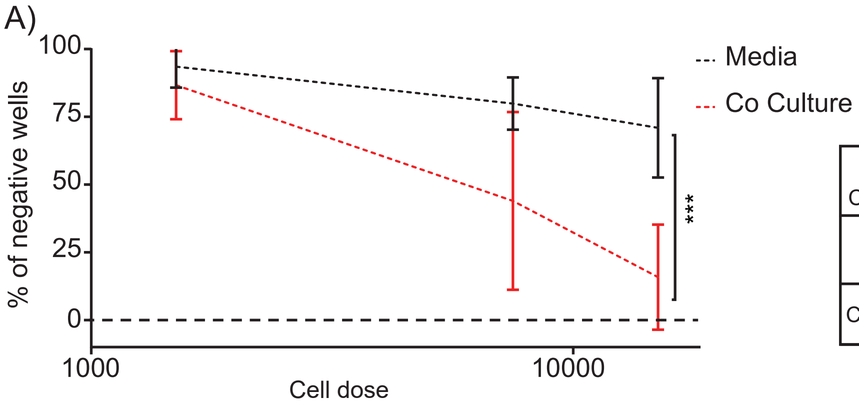


| Culture Conditions | IR dose (Gy) | Frequency | IC 95% |
|---|---|---|---|
| Medium | 0 | 1/102 | 1/127 ; 1/81.2 |
| | 3 | 1/35685 | 1/46,864 ; 1/27,173 |
| Co-Culture | 0 | 1/127 | 1/157 ; 1/103 |
| | 3 | 1/7446 | 1/9501 ; 1/5835 |

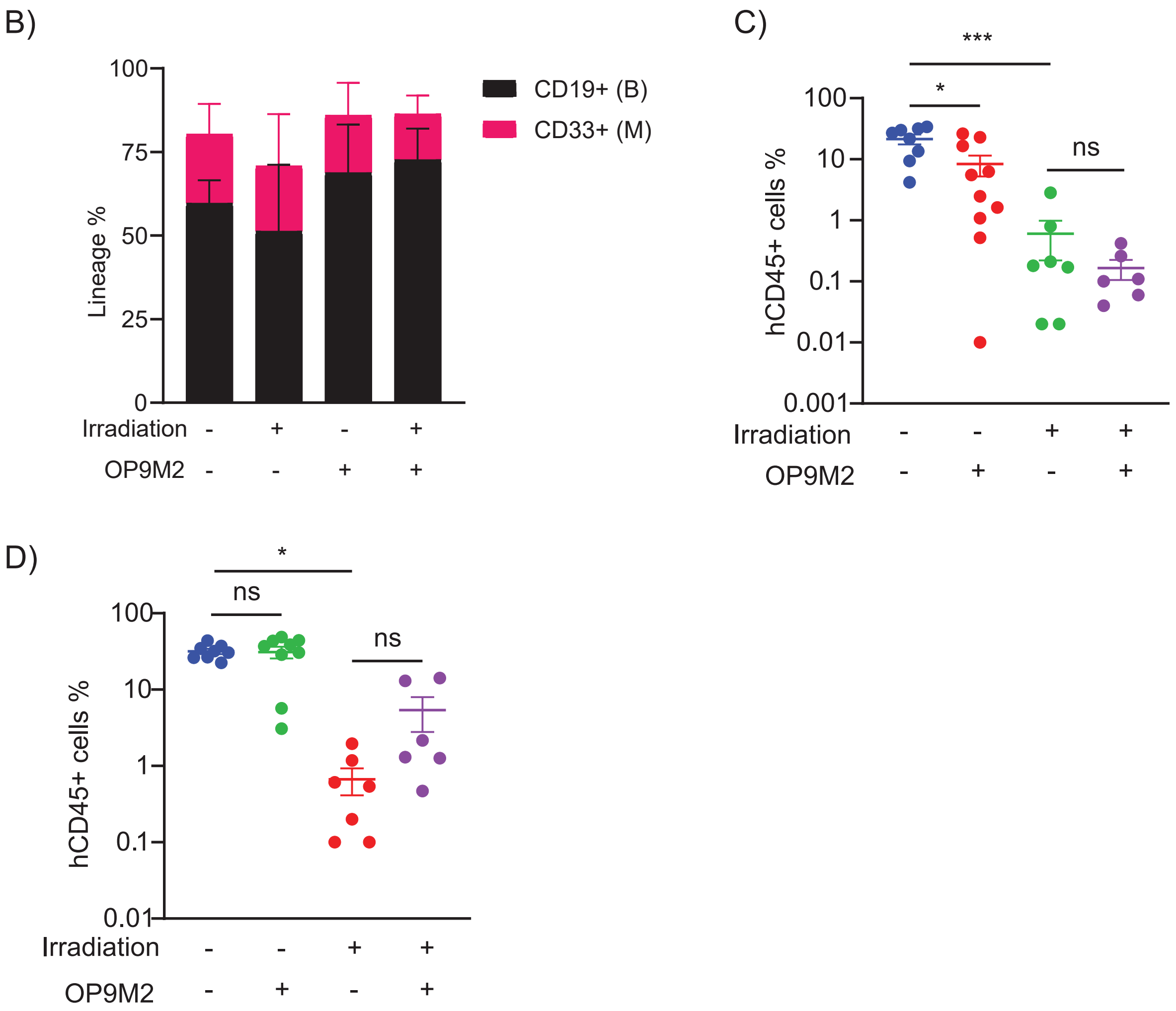

Supplemental Figure 4

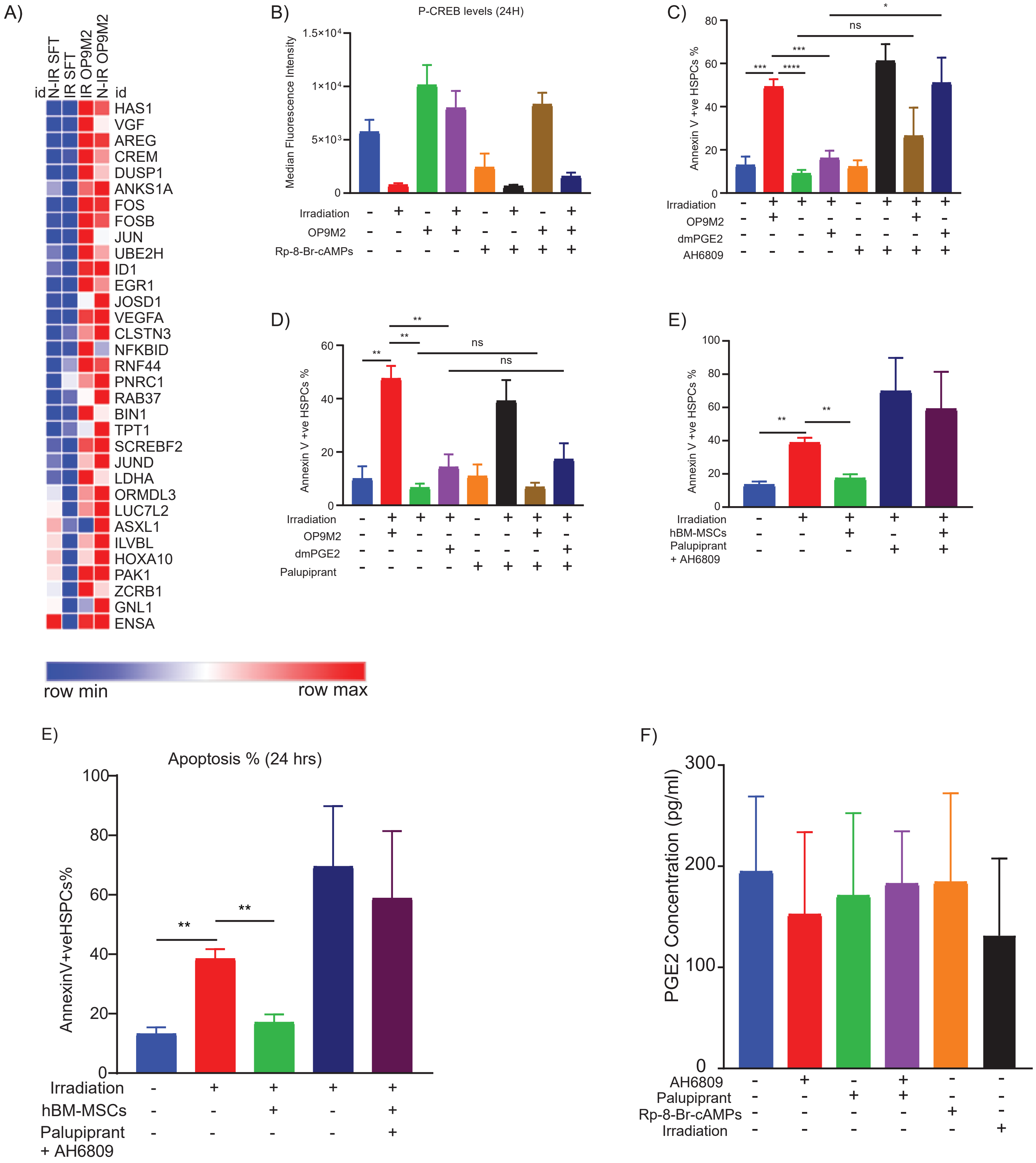

Supplemental Figure 5

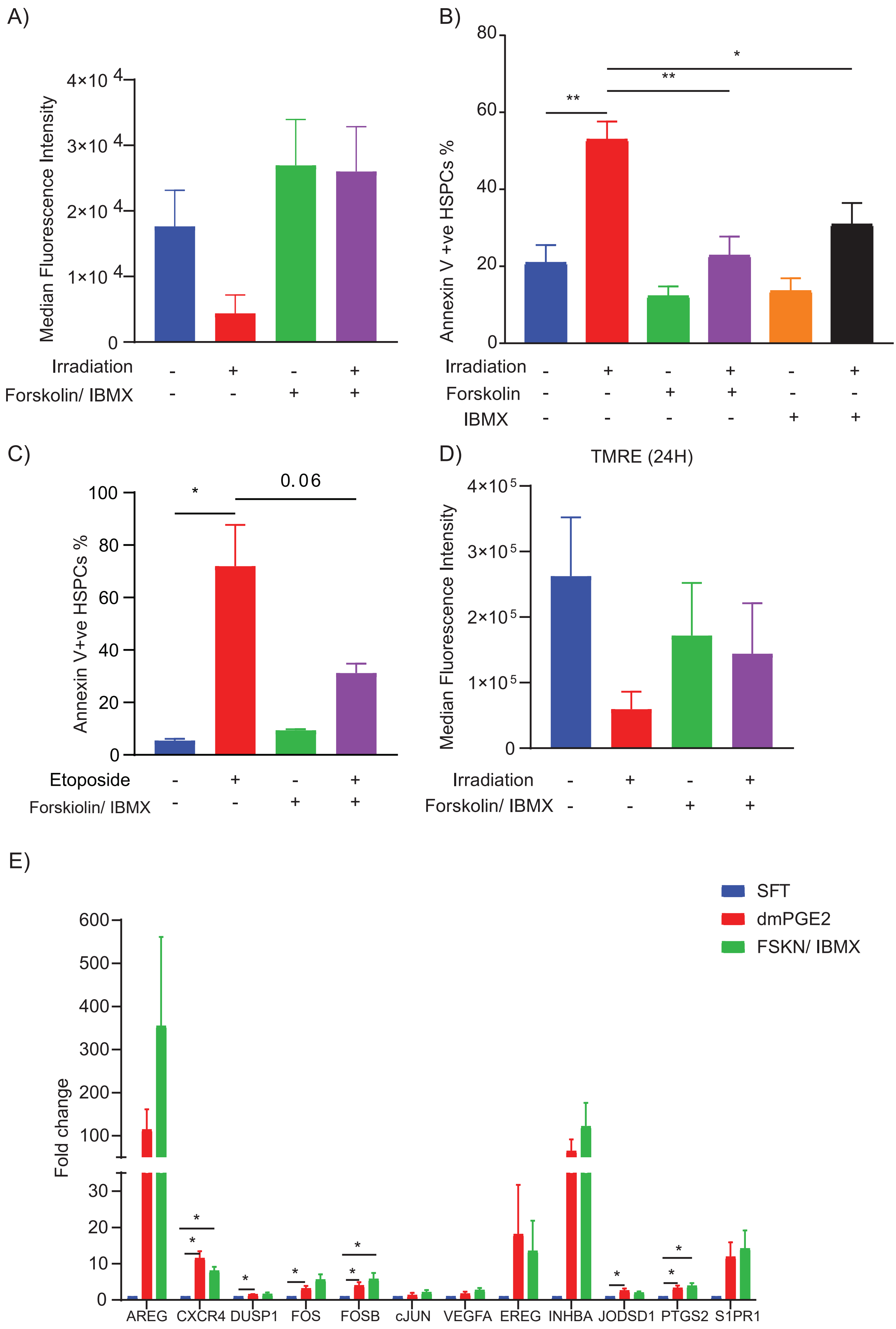

Supplemental Figure 6

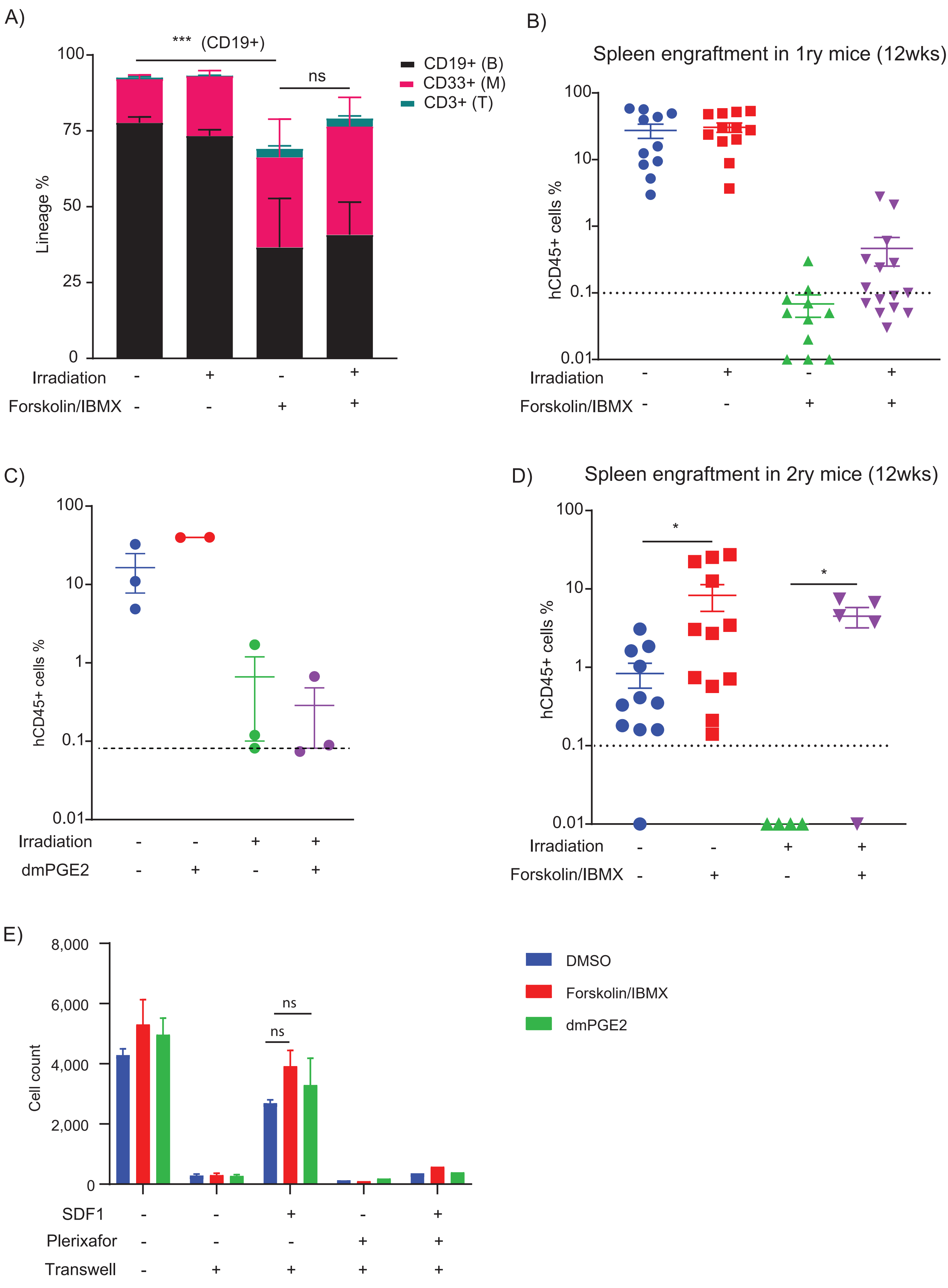

Supplemental Figure 7

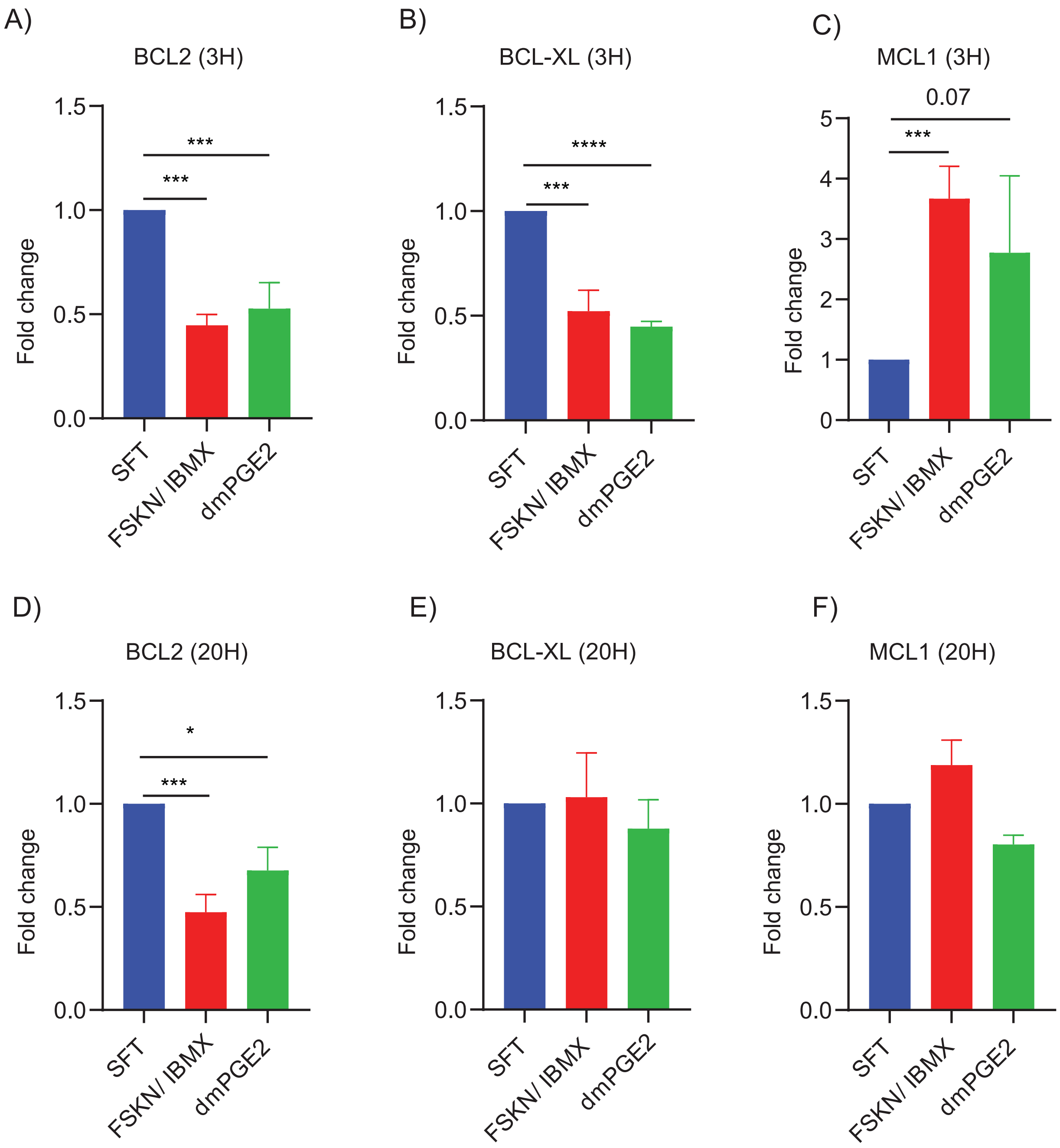

Supplemental Figure 8

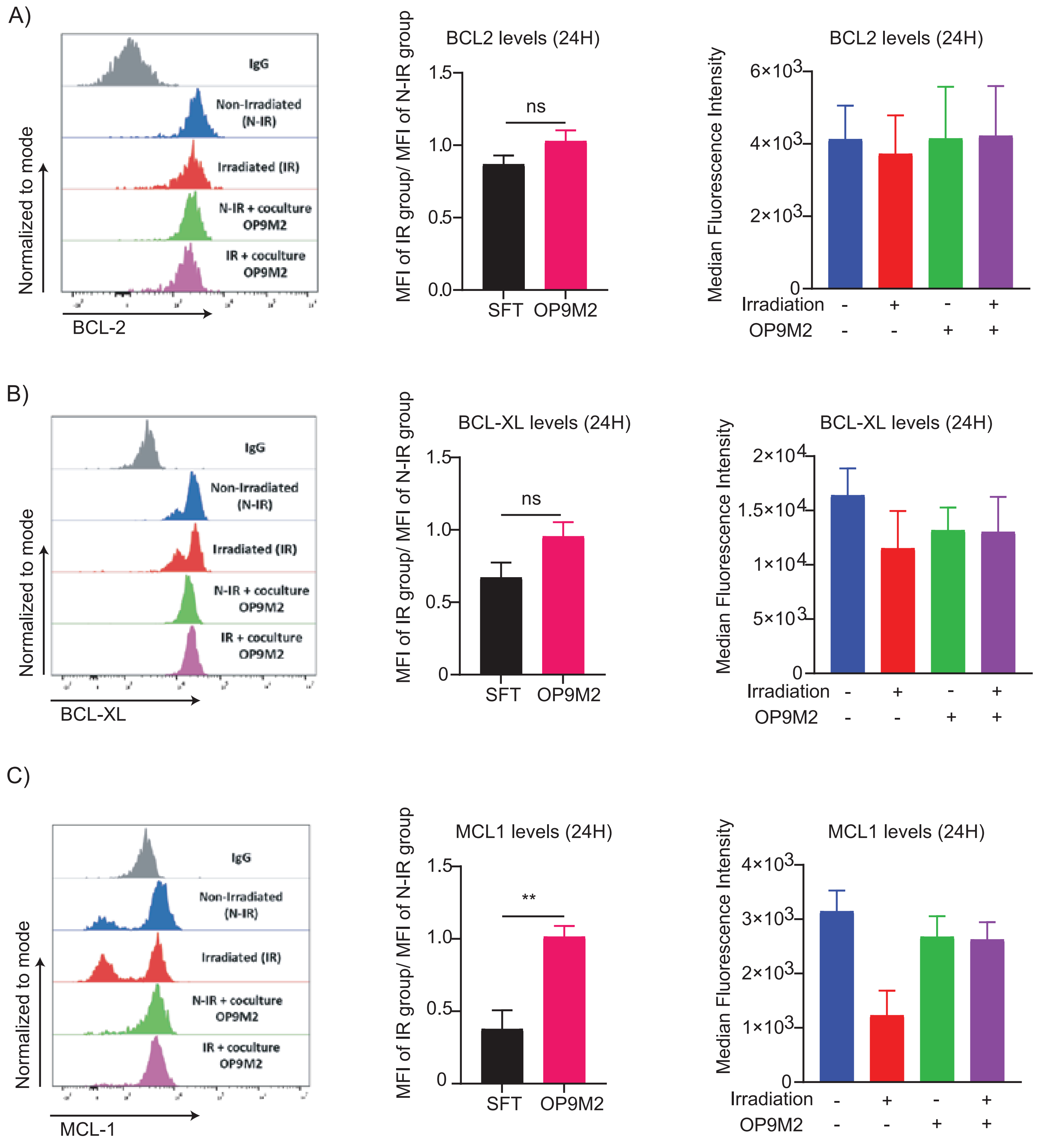

Supplemental Figure 9

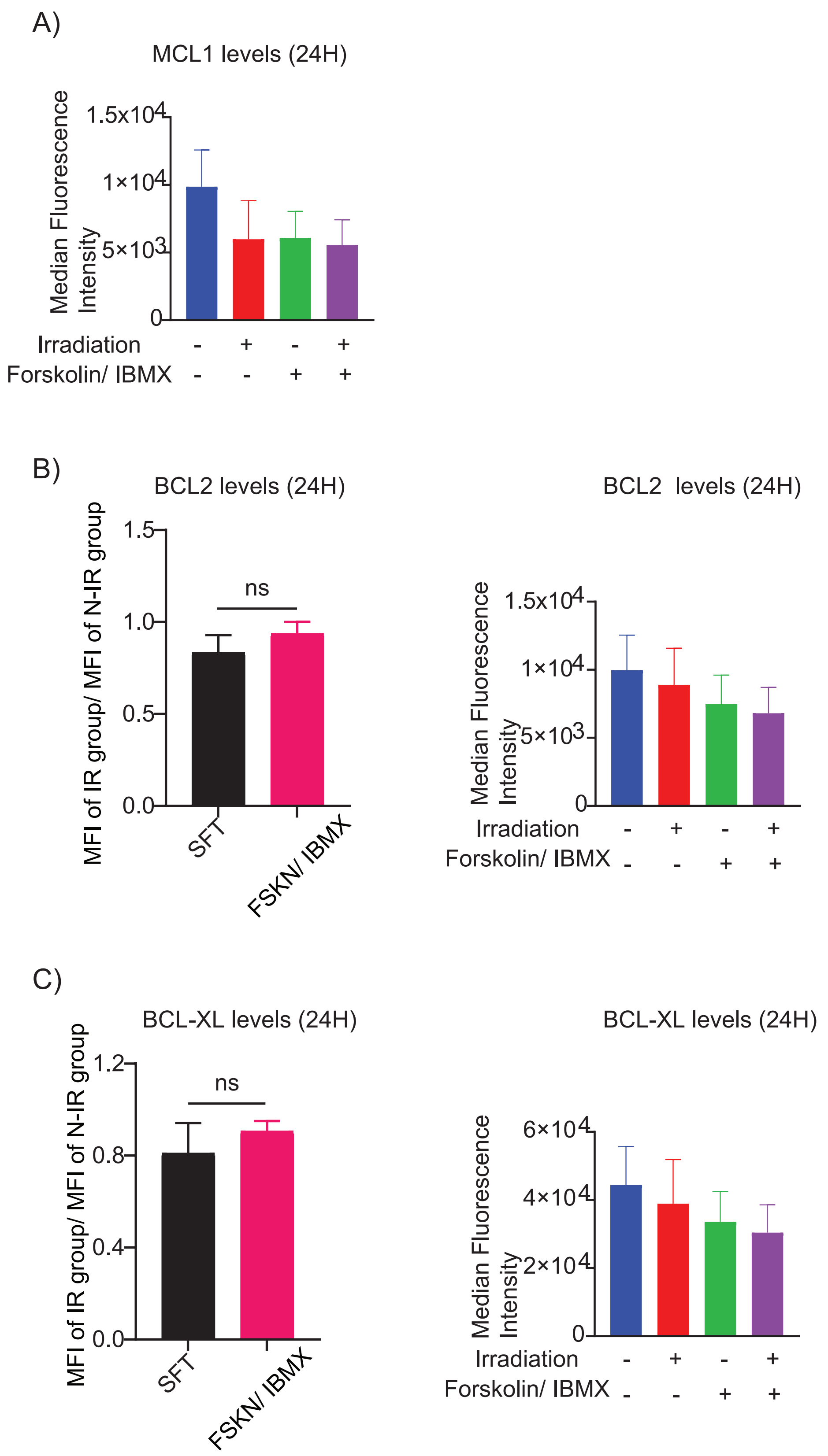

Supplemental Figure 10

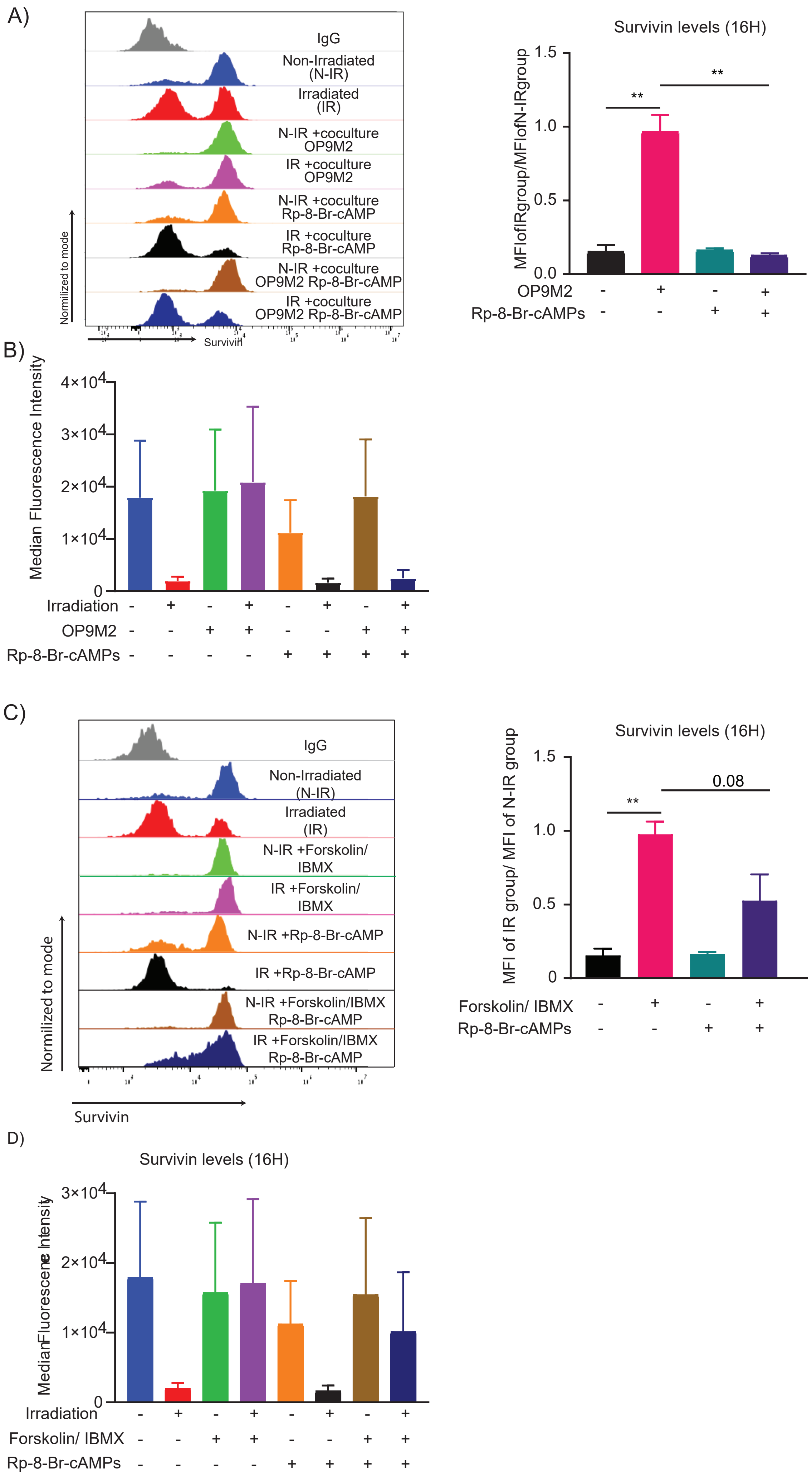
